# Metabolic Response of a Chemolithoautotrophic Archaeon to Carbon Limitation

**DOI:** 10.1101/2025.02.13.638070

**Authors:** Logan H. Hodgskiss, Melina Kerou, Zhen-Hao Luo, Barbara Bayer, Andreas Maier, Wolfram Weckwerth, Thomas Naegele, Christa Schleper

## Abstract

The ubiquitously distributed ammonia-oxidizing archaea generate energy from ammonia and build cell mass from inorganic carbon sources, thereby contributing to both the global nitrogen and carbon cycles. However, little is known about the regulation of their predicted core carbon metabolism. A thermodynamic model for *Nitrososphaera viennensis* was developed to estimate the consumption of inorganic carbon in relation to ammonia consumed for energy and was tested experimentally by growing cells in carbon-limited and excess conditions. A combined proteomic and metabolomic approach of the experimental conditions revealed distinct metabolic adaptation depending on the amount of carbon supplied, either in a catalase or pyruvate background as a reactive oxygen species scavenger. Integration of protein and metabolite dynamics revealed a cellular strategy under carbon limitation to maintain a pool of amino acids and an upregulation of proteins necessary for translation initiation to stay primed for protein synthesis. The combination of modelling and functional genomics fills gaps in the understanding of the central metabolism and its regulation in a chemolithoautotrophic, ammonia-oxidizing archaeon even in the absence of available genetic tools.

## Introduction

Ammonia-oxidizing archaea (AOA) have been found in a wide variety of environments often outnumbering their bacterial counterparts, ammonia-oxidizing bacteria (AOB)^1–7^. Based on their abundance, their contributions to the global nitrogen cycle are substantial. While there are many efforts to unravel the archaeal ammonia oxidation pathway^8–13^, their carbon metabolism also draws attention as all so far characterized AOA are autotrophic and therefore represent a direct biological link between the global nitrogen and carbon cycles.

AOA participate in fixation of inorganic carbon by a unique version of the 3-hydroxpropionate/4-hydroxybutyrate cycle. Although it appears similar to the 3-hydroxypropionate/4-hydroxybutrate cycle in hyperthermophilic archaea^14^, it is more efficient due to the use of only one ATP equivalent (rather than two) in selected enzymatic reactions and the use of promiscuous enzymes that can catalyze multiple steps in the pathway^14,15^. Most of the biochemical work on the AOA carbon fixation cycle has been carried out with the first isolated AOA, *Nitrosopumilus maritimus*^15–17^, which was isolated from a marine aquarium and represents the family *Nitrosopumilaceae*^18^ (GTDB taxonomy^19^, used throughout). Since then, genome comparisons have found this cycle to also be present in the first soil isolate, *Nitrosospheara viennensis*^20^, and to be conserved in all AOA^11,21^.

While the carbon fixation pathway within AOA has been identified, the regulation of this pathway is not well characterized. The tricarboxylic acid (TCA) cycle, gluconeogenesis, and non-oxidative pentose phosphate pathway, have all been identified as conserved central carbon pathways^11,21^, but the response and control of these pathways remains unclear. Considering that terrestrial organisms like *N. viennensis* experience strong fluctuations of their growth substrates and variations in environmental parameters, investigations of limiting conditions by functional genomics techniques can give insights into the adaptive capacity of the organisms. To this end, inorganic carbon was chosen as a limiting substrate in this study. Unlike nitrogen and oxygen, inorganic carbon is not directly tied to the amount of energy available to AOA. It also allows to investigate what pathways and proteins respond within the central carbon metabolism to maintain a flow of carbon, shedding light on starvation responses, resource allocation choices, and interconnections with other environmental response pathways which may highlight other physiological points of interest within the cell.

Knowing the primary energetic reactions that drive AOA metabolism (ammonia oxidation and oxygen reduction), an energetic model was calculated in this study based on the Thermodynamic Electron Equivalents Model 2 (TEEM2)^22^ method to estimate the amount of inorganic carbon the organism would need to effectively grow. Once calculated, *N. viennensis* was grown at five different carbon concentrations with the addition of either catalase or pyruvate to test the resulting model. Proteins and metabolites were simultaneously evaluated via a combined extraction protocol and analyzed in a systems biology framework to better understand how *N. viennensis* metabolically behaves under limited carbon supply. The results show distinct shifts in the proteome and metabolome and highlight unique and distinct roles of multiple oxidative stress proteins within the cell. These results contribute to a more complete picture of redox control and the central carbon metabolism of AOA.

## Results

### Growth of N. viennensis agrees with thermodynamic predictions

A thermodynamic model of the ammonia-oxidizing archaeon *Nitrososphaera viennensis* predicted a carbon consumption of 0.064 mols of inorganic carbon per 1 mol of ammonia (0.128 mol carbon/2 mol ammonia) consumed with an overall growth equation:

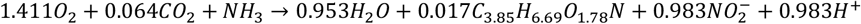

The model takes into account physiological traits of ammonia-oxidizing archaea including a monooxygenase reaction, estimated biomass electron equivalents, activation of carbon to acetyl-CoA, and an assumed energy transfer efficiency (see Materials and Methods)^22,23^. *N. viennensis* was grown at varying carbon concentrations (Figure 1A) in a closed system using either exogenous catalase or pyruvate to diminish the effect of reactive oxygen species (ROS)^24^ (Figure 1B,C) while using nitrite production as a proxy for growth.

**Figure 1:**
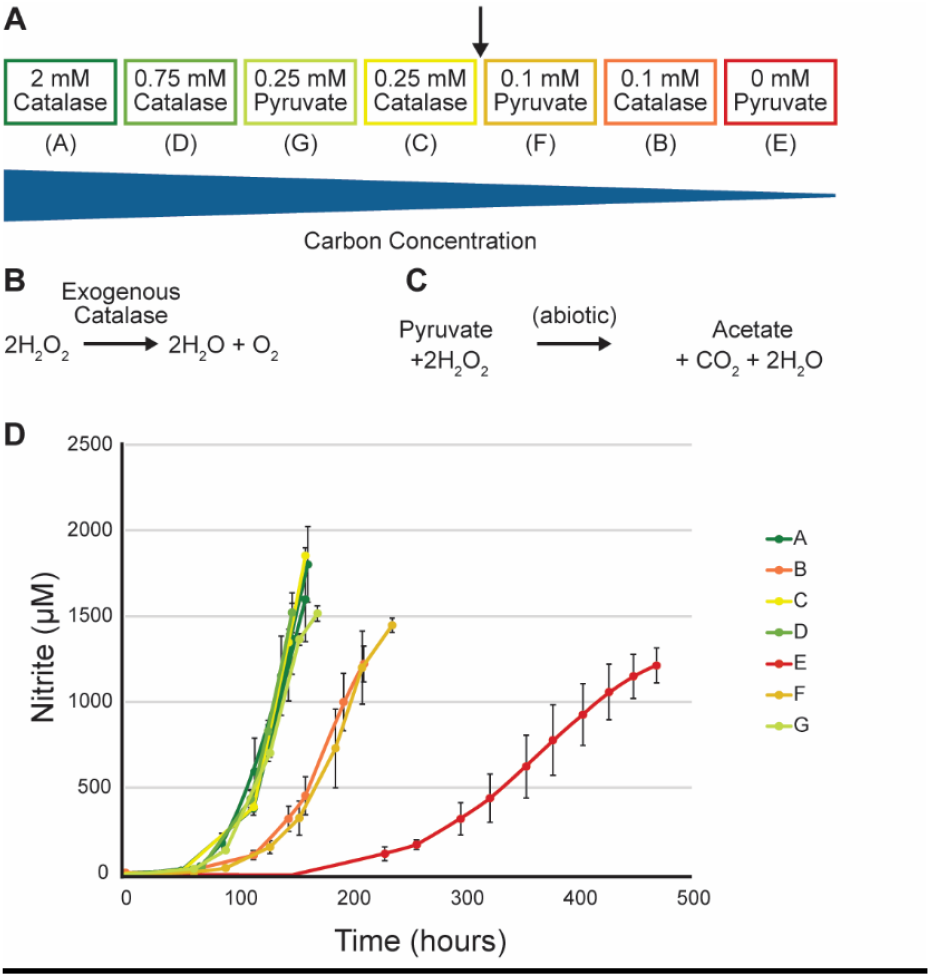
Growth curves of *N. viennensis* with varying amounts of inorganic carbon. **A.)** Experimental design of different inorganic carbon concentrations. **B.)** Reaction of catalase with hydrogen peroxide. Black arrow indicates the calculated, theoretical carbon threshold of 0.12 mM with respect to a 2 mM ammonia concentration. **C.)** Reaction of pyruvate with hydrogen peroxide. **D.)** Growth curves of *N. viennensis* exhibiting responses to varying levels of inorganic carbon.

All cultures were inoculated with comparable amounts of cells but started with carbon concentrations ranging from 0 mM – 2 mM sodium bicarbonate and the same amount of ammonia (2mM). Cultures incubated with inorganic carbon above the theoretical threshold of 0.128 mM grew without obvious limitations (Figure 1D; A,C,D,G). Cultures grown below this threshold were either slightly limited (0.1 mM carbon; B,F) or extremely limited (0 mM carbon; E). As ammonium was present in the medium, limited growth was directly attributed to lack of carbon rather than an energy limitation (Figure EV1). Cultures were harvested in late exponential phase when possible. Condition B (0.1 mM carbon with catalase) was harvested earlier than other cultures before it reached stationary phase due to a lack of carbon. The most limited condition (E) with 0 mM of supplied inorganic carbon presented cultures with an extended lag phase and very slow growth as measured by nitrite production. In these cultures, the only available inorganic carbon came from added pyruvate that was decarboxylated in the presence of hydrogen peroxide (Fig. 1C). The use of pyruvate or catalase as a ROS scavenger also produced a notable difference in the other limited cultures. All cultures of condition F and condition B were grown with 0.1 mM inorganic carbon. However, the use of pyruvate in condition F to counter ROS provided an additional release of CO_2_. This slight difference resulted in F cultures being able to oxidize more ammonia than condition B, which entered stationary phase at ∼1200 μM NO_2_^-^ (Figure EV2).

Carbon balances were calculated using initial and final carbon concentrations in the aqueous and gas phases through dissolved inorganic carbon measurements and gas chromatography respectively (see Supp. Materials and Methods). Differences in carbon consumption were observed even among cultures with similar growth curves (Figure EV3). Conditions with an excessive amount of carbon (conditions A and D; 2 mM and 0.75 mM, respectively) consumed a higher amount of carbon/nitrite produced (mol/mol) than cultures that were closer to the theoretical carbon threshold (conditions C and G) or under the threshold (conditions B, F, and E) (Figure EV3). The higher amount of consumed carbon could have been incorporated into the carbon storage molecule polyhydroxybutyrate (PHB) or alternatively formed precipitates that would not be captured in the inorganic carbon measurements. Protein and DNA levels normalized to nitrite produced in cultures of the extremely carbon-limited condition E indicate a lower number of cells and a decoupling of ammonia oxidation and cell growth in extremely limited cultures (Figure EV3).

### Differential protein expression in carbon-limited conditions reveals the main anabolic routes within the cell

Proteomes of the seven different conditions were subjected to a principal component analysis (PCA) (Figure 2). 1264 proteins of 3,123 predicted protein coding genes (40.47%) were identified from the combined data. All conditions above the carbon threshold grouped together on the left side of principal component one (PC1) while the carbon limited cultures grouped on the right side of PC1. Condition E, representing the most limited culture, grouped the farthest from carbon replete cultures with conditions B and F, representing slightly limited cultures, were found between these two groupings. The separation of proteomes on the PCA plot closely reflects the observed growth behavior (Figure 1).

**Figure 2:**
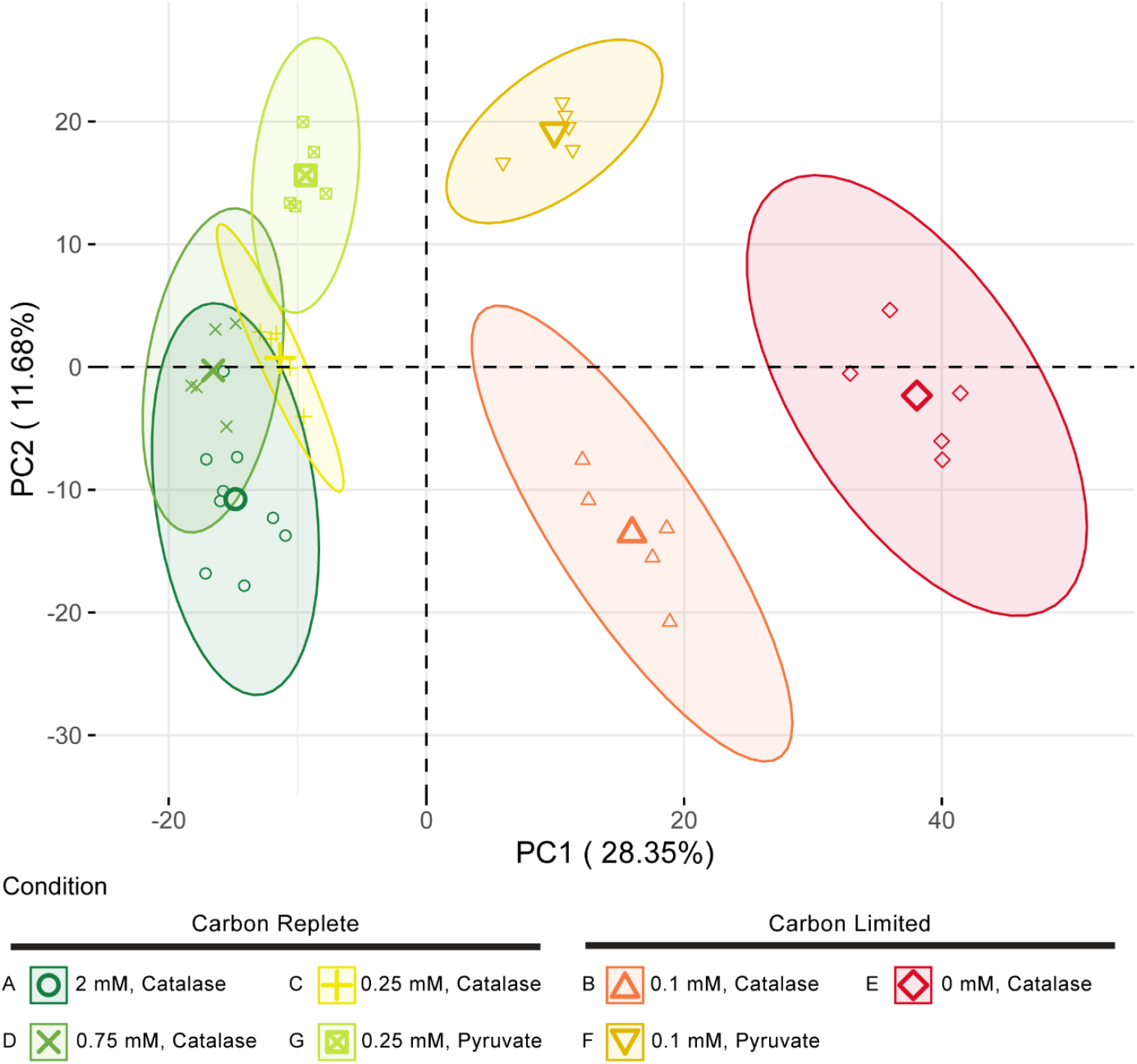
Principal component analysis of *N. viennensis* proteomes. Variance within the proteomes is best described by principal component 1 (PC1) with non-limited conditions A, D, C, and G on the far left of PC1 and the extremely limited culture on the far right. Limited cultures B and F fall between these two groupings on PC1. Ellipses represent confidence ellipses at a level of 0.95. Centers of ellipses are marked by bold points within each ellipse respectfully.

Although the PCA showed overall differences among the proteomes of the conditions, the most abundant proteins for each condition were largely the same. Of the most abundant proteins (126 proteins, 10% cut-off) from each condition, 100 were shared (Dataset EV1). Those included mostly proteins involved in energy and carbon metabolism as well as translation (i.e., ribosomal proteins). The high relative abundance of these proteins across all conditions indicate that they are functionally important for the cell regardless of the limitation of carbon resources.

Following the principal component analysis, clustering was performed on the proteomic data to identify specific proteins associated with the various conditions and statistical tests (either ANOVA or Kruskal-Wallis) were used to determine which proteins showed a statistical difference (adjusted *P* value ≤ 0.05) among the tested conditions. Approximately 80.7% of the total detected proteins showed a change among conditions with 20.3% remaining constant.

Proteins were divided into 7 clusters after hierarchal clustering (Figure 3A). The carbon limited cluster (Figure 3A, Cluster V) represents proteins that increased in relative abundance under extreme carbon limitation. A majority of proteins (15 out of 17) making up the 3-hydroxypropionate/4-hydroxybutyrate carbon fixation cycle was found in this cluster. Based on archaeal clusters of orthologous genes (arCOG) categories^25,26^ and a hypergeometric test of all detected proteins, the E cluster was also enriched for proteins involved in amino acid transport and metabolism (Figure 3B). In contrast, the carbon replete cluster (Figure 3A, Cluster VII) representing high relative abundance in cultures that were not limited (conditions A, C, D, and G) showed enrichment for cell cycle control, transcription, and translation (Figure 3B). These included proteins found in the Cluster VII included many ribosomal proteins as well as cell division proteins from the CDV system found to function in AOA^27^.

**Figure 3:**
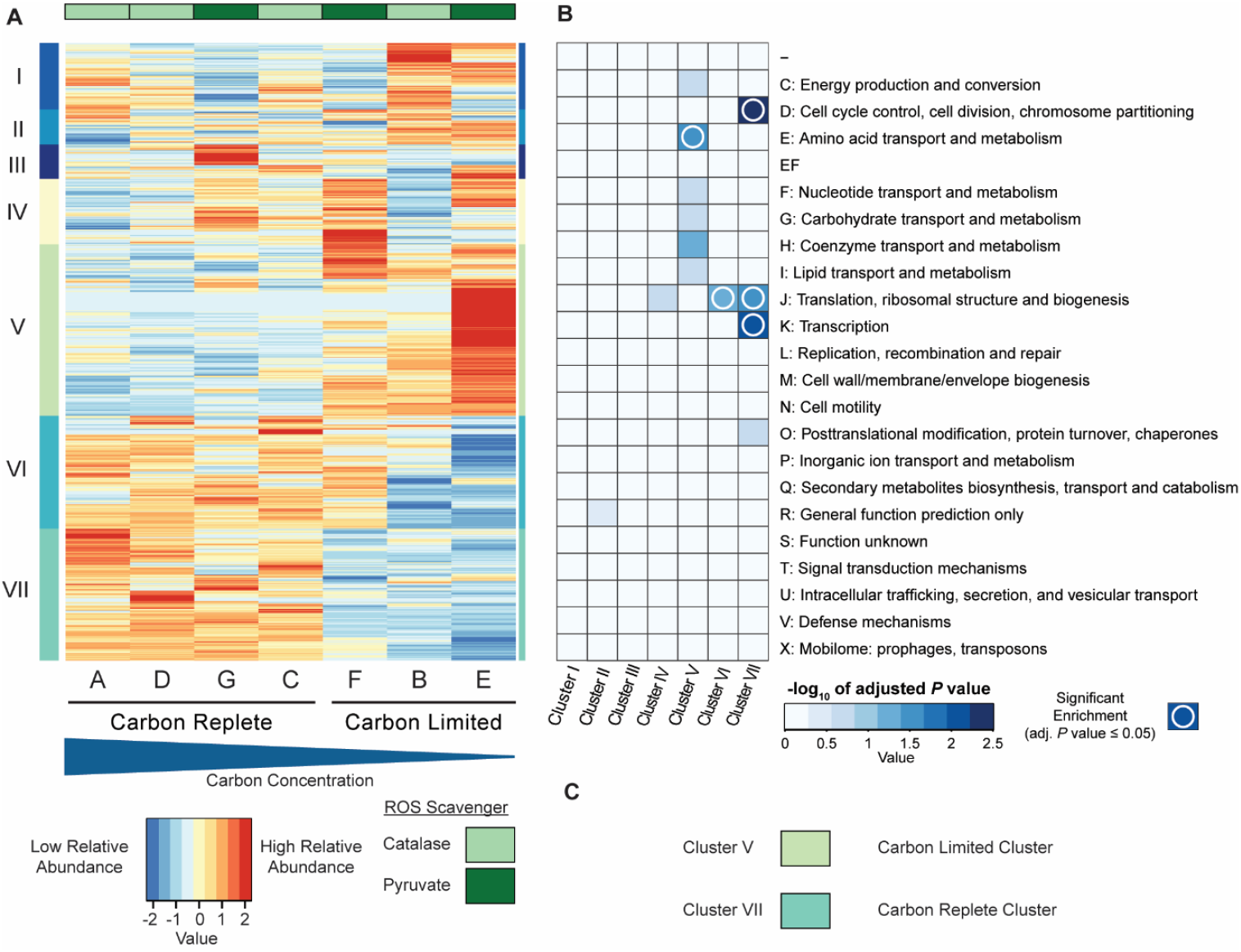
Heatmap and enrichment of hierarchically clustered proteins. **A.)** Proteins were clustered after being centered and scaled. Clustered proteins were divided into 7 clusters that are represented by the different colors and Roman numerals on the left side of the heatmap. Each horizontal line represents the relative abundance of a specific protein across all conditions. Conditions are organized from highest carbon concentration to lowest carbon concentration from left to right. Color bars across the top represent conditions with either catalase (light green) or pyruvate (dark green) as a reactive oxygen species scavenger. **B.)** Protein archaeal clusters of orthologous genes (arCOGs) enrichment analysis. Boxes with a white circle indicate arCOG categories that are enriched in their respective cluster based off of a hypergeometric test. EF represents a combined category of amino transport and metabolism (E) and nucleotide transport and metabolism (F). **C.)** Clusters of interest with respect to carbon limitation.

Within proteins of the 3-hydroxypropionate/4-hydroxybutyrate carbon fixation cycle, statistical tests showed that the average relative abundance (reported as LFQ values) for these proteins under carbon limitation (0 mM; condition E) were significantly different when compared to the standard carbon concentration (2mM; condition A). A closer look at the primary carboxylating protein, acetyl-CoA carboxylase, revealed a strong reaction in the relative abundance of this protein across all conditions (increased abundance with less inorganic carbon), even if an effect was not observed in the growth curves (Figure EV4). In the case of the so far unknown proteins for the reduction of acryloyl-CoA and succinic semialdehyde, 10 candidate proteins had been proposed earlier based on a genomic analysis^11^. The clustering analysis presented here would suggest that Adh4 (NVIE_024420) is the protein fulfilling one or both of these roles in the cycle. As expected, proteins participating in the production (PhaE, PhaC) of the potential storage compound polyhydroxybutyrate (PHB) were shown to markedly decrease in the proteome under carbon limiting conditions while proteins that would funnel carbon away from PHB and towards the replenishment of acetyl-CoA (PhaA, PhaB) were increased under carbon limitation (Figure 4). The only two carbon fixation cycle proteins not found in this cluster were methylmalonyl-CoA epimerase, which decreased under carbon limitation, and hydroxyacyl-CoA hydrogenase which showed some variation among conditions containing 0.1-0.25 mM of inorganic carbon rather than a strong response under limitation.

**Figure 4:**
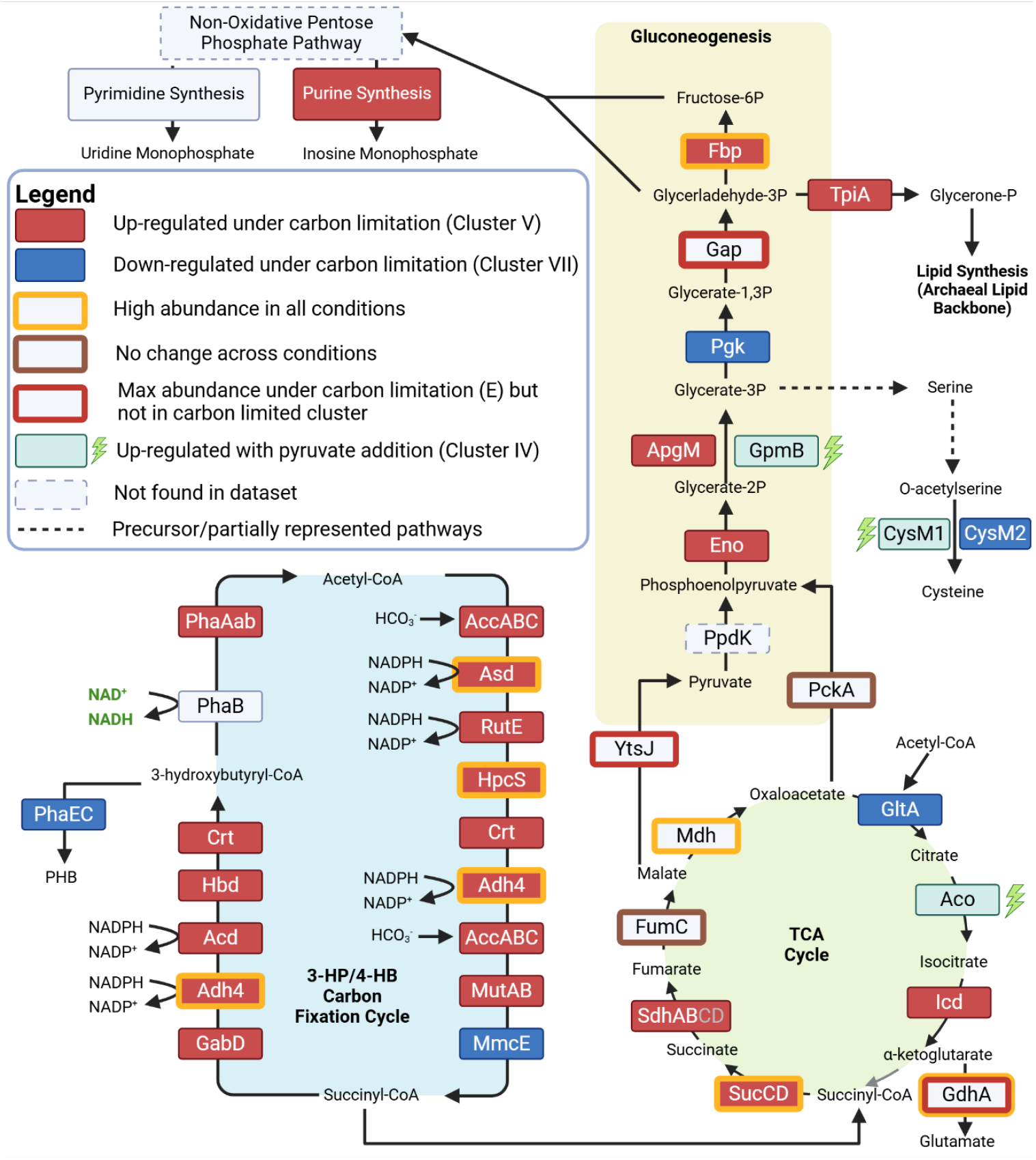
*Nitrososphaera viennensis* core metabolism under carbon limitation. Protein abbreviations are found in rectangular boxes. Up-regulated protein under carbon limitation is defined as proteins found within the carbon limited cluster (Cluster V) that show a statistical difference between condition E and condition A. Down-regulated proteins under carbon limitation are defined as proteins found in the carbon replete cluster (Cluster VII) that show a statistical difference between condition E and condition A. Proteins upregulated with addition of pyruvate were found in Cluster (IV) and showed a statistical difference between at least two conditions. Locus tags and accession number for proteins can be found in Dataset EV1. Created in BioRender. Hodgskiss, L. (2025) https://BioRender.com/l00t241.

Other proteins within this cluster were found in the TCA cycle, gluconeogenesis, pentose phosphate pathway, and acetyl-CoA biogenesis. Within the TCA cycle, proteins following the putative incorporation of succinyl-A were up-regulated including succinyl-CoA ligase (SucD, SucC) and subunits of succinate dehydrogenase (SdhA, SdhB). Conversely, the subsequent step of acetyl-CoA incorporation, citrate synthase, was down-regulated under carbon limitation (Figure 4). Fumarate hydratase exhibited no change. At the branching point of malate, malate dehydrogenase (leading to oxaloacetate) was down-regulated (Cluster VI) while malic enzyme (leading to pyruvate) was up-regulated (Cluster III). Regardless, malate dehydrogenase maintained a higher relative abundance than malic enzyme across all conditions.

Other proteins of interest included five of six P-II regulatory proteins that were also detected with three (CnrA, NVIE_003920; CnrD, NVIE_014570; and CnrC, NVIE_014550) showing an increase under carbon limitation (Cluster V) and two (GlnB NVIE_007790, Cluster VI; and CnrB, NVIE_013340, Cluster VII) showing a decrease under carbon limitation.

Additionally, in a supervised partial least squares discriminant (PLS-DA) analysis, protein NVIE_010650 was found to be highly responsive to carbon limitation (Figure EV5). While no functional BLAST hits were obtained, a structural search revealed that the protein contains a putative photosynthetic reaction center (PRC) domain related to potential ribosome maturation proteins.

### Metabolomic analysis reveals an accumulation of amino acids under carbon limitation

Selected metabolites were identified and quantified based on standard curves produced from known stock concentrations. Unfortunately, a few of the organic acids in samples could not be accurately determined due to contamination from the highly purified water system as determined by comparison to background tests. Therefore, these metabolites (citric acid, lactic acid, and oxaloacetate) had to be excluded from the analysis. Pyruvate was also excluded due to its addition to conditions E, F, and G as a ROS scavenger. Metabolites that could be accurately quantified were normalized to the total amount of carbon consumed for each culture and were clustered to identify metabolites associated with specific conditions. The normalization would therefore represent metabolites in terms of carbon-moles of metabolite produced per mole of carbon consumed (see Materials and Methods). A metabolite PCA plot showed a clear separation of the most limited condition (E) (Figure EV6). In a clustering analysis, almost all amino acids were clustered with the most limiting carbon condition, condition E (Figure 5). Identifiable sugars (glucose, maltose, and trehalose), were almost exclusively associated with conditions that used catalase as a ROS scavenger (A,B,C,D, Figure EV7). Melibiose, a disaccharide, was an exception to this trend and was found predominantly in conditions that were carbon limited (Figure EV7). Fumaric acid, galactose, and gluconic acid had a higher relative abundance within cultures that were not carbon limited.

**Figure 5:**
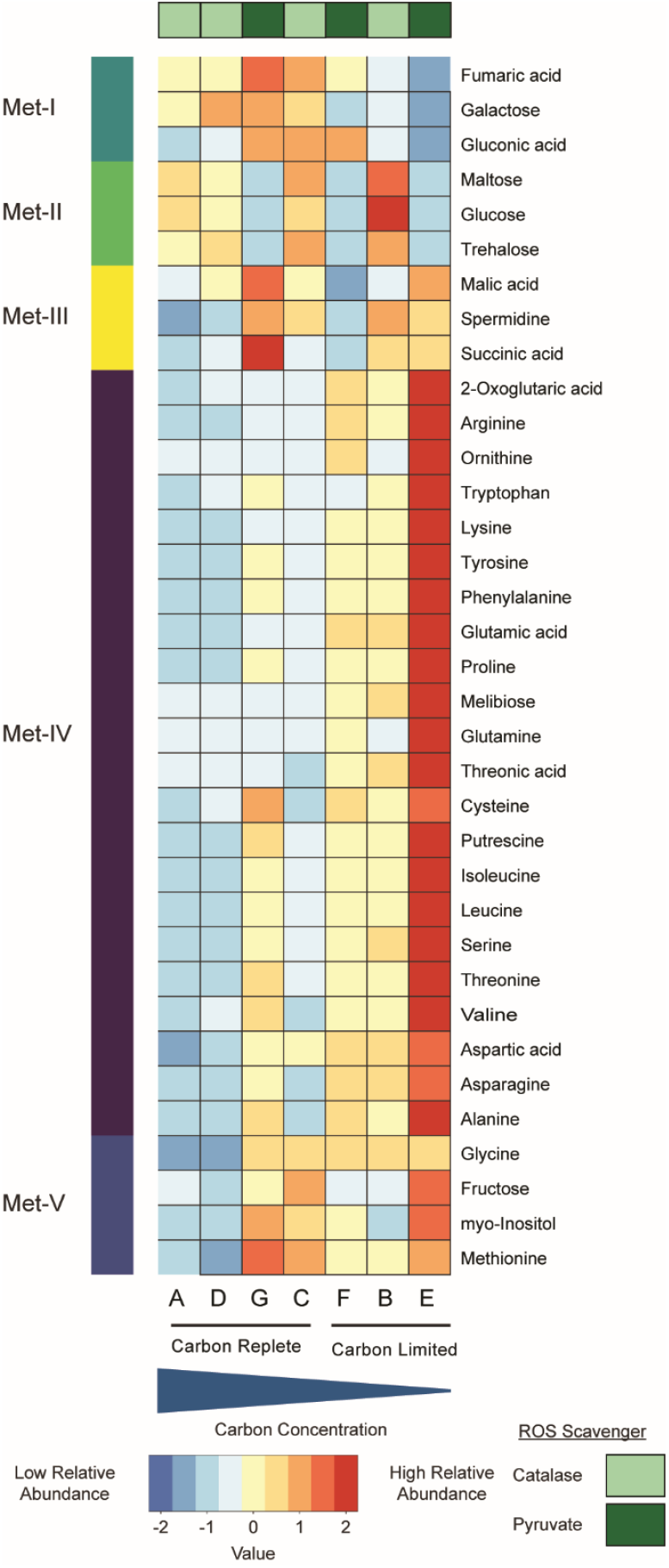
Heatmap of hierarchically clustered metabolites. Metabolites were clustered after being centered and scaled. Clustered metabolites were divided into 5 clusters that are represented by the different colors on the left side of the heatmap. Each horizontal line represents the relative abundance of a specific metabolite across all conditions. Conditions are organized from highest carbon concentration to lowest carbon concentration from left to right. Color bars across the top represent conditions with either catalase (light green) or pyruvate (dark green) as a reactive oxygen species scavenger.

### Combined analysis of proteins and metabolites identifies key cellular processes when exposed to environmental stress

Proteomic and metabolomic data were analyzed together to further investigate the metabolism of *N. viennensis*. Metabolites identified to be relatively abundant in condition E (Cluster Met-IV) were analyzed with all proteins across all conditions to identify significant correlations (i.e., concurrent pattern of proteins increasing with metabolites). This revealed several proteins that consistently correlated with multiple metabolites. Within these correlations, 33 proteins were identified that correlated with 5 or more metabolites (adjusted *P* value ≤ 0.05, R ≥ 0.75). The four proteins with identified functions that most often correlated with metabolites were methionine sulfoxide reductase A (MsrA) with 15 metabolite correlations, and acetylornithine aminotransferase (ArgD2), a translation initiation factor (eIf-2a), and inosine-5’-monophosphate dehydrogenase (GuaB) each with 9 correlations (Dataset EV1). Eighteen of the 33 proteins had predicted functions, three of which were involved in translation (arCOG category J).

The relationship between MsrA and metabolites was further explored by correlating MsrA with all other proteins to find similar behavioral patterns. MsrA had positive correlations (adjusted *P* value ≤ 0.5, R ≥ 0.7) with 59 proteins (Dataset EV1). Several proteins that correlated with MsrA also had multiple metabolites (>=5). This included two of the three translationally involved proteins: the translation initiation factor eIf-2a and methionine aminopeptidase (Map) (Dataset EV1).

MsrB2, another methionine sulfoxide reductase found in the proteome exhibited a very different pattern compared to MsrA (Figure EV8). It was not increased under carbon limitation, but was expressed significantly higher in all conditions that were supplemented with pyruvate as a ROS scavenger rather than catalase.

A correlation analysis (adjusted *P* value ≤ 0.5, R ≥ 0.75) was also performed on all proteins with the sugars trehalose, maltose, and glucose which only appeared in conditions with catalase (Figure EV7). Five proteins were found to be correlated with all three sugars. All five had positive correlations and included ABC efflux transporter proteins (NVIE_028580 and NVIE_028560), a putative thiamine biosynthesis protein (NVIE_000430), a probable zinc uptake transporter (NVIE_021020), and a putative transcriptional regulator (NVIE_002180).

A correlation analysis (adjusted *P* value ≤ 0.5, R ≥ 0.75) with melibiose, which only appeared under carbon limitation, revealed a wide range of correlations with 72 positive and 32 negative protein correlations (Dataset EV1).

### Carbon metabolism and detoxification proteins responding to ROS scavenger

Due to the response of MsrB2 to the choice of ROS scavenger, other known ROS detoxification proteins and core carbon metabolism proteins were manually checked for differences between the choice of ROS scavenger rather than carbon limitation. Multiple oxygen detoxification proteins and some carbon metabolism proteins were also found with MsrB2 in a protein cluster that responded specifically to the addition of pyruvate (Cluster IV). In addition to MsrB2, these included an alkyl hydroperoxide reductase (Ahp2, NVIE_011770), the 2,3-bisphosphateglycerate-dependent phosphoglycerate mutase (GpmB) of the gluconeogenesis pathway, and the mitochondrial-type aconitase, Aco, albeit with less differences between conditions based on a post hoc Tukey analysis (Figure EV9). MsrA and TrxB were the only detoxification proteins found to increase under carbon limitation (Cluster V). Oxidative stress detoxification proteins found to decrease under carbon limitation (Cluster VII) included an alkyl hydroperoxide reductase (Ahp1, NVIE_013750), superoxide dismutase (SodA, NVIE_030260), and a putative thioredoxin (TrxX, NVIE_024000). Although decreasing under carbon limitation, SodA and Ahp1 were included with proteins that exhibited the highest relative abundance across all conditions. Two thioredoxins (TrxA1 and TrxA2, NVIE_029260 and NVIE_030000 respectively, both in Cluster VI) were also found to decrease under carbon limitation.

## Discussion

### Thermodynamic model based on primary metabolic reaction accurately predicts carbon consumption in ammonia oxidizers

The experimental design for carbon limitation of *Nitrososphaera viennensis* was primarily guided by a thermodynamic model based on the Gibbs free energy of the electron acceptor and electron donor reactions of ammonia oxidizers. This model was able to predict the needed amount of carbon for a respective amount of nitrogen (represented as ammonia) that is oxidized (0.06 mol C/mol N) and was used to establish starting inorganic carbon concentrations both above and below a calculated threshold. The use of pyruvate as a ROS scavenger in selected conditions (G, F, and E) allowed for an added boost of inorganic carbon as it was abiotically decarboxylated by hydrogen peroxide produced by the cells during growth.

The result of the model (0.06 mol C/mol N) falls directly between observed carbon fixation yields of marine ammonia-oxidizing archaea (0.1 mol C/mol N) and bacteria (0.045 mol C/mol N) using radioisotope measurements from Bayer et al.^28^, supporting the validity of the assumptions made when creating the model (see Materials and Methods). The differences in observed carbon fixation rates between the two nitrifying clades can be attributed to the different carbon fixation pathway used by each clade with AOA utilizing the more efficient 3-hydroxypropionate/4-hydroxybutyrate( 3-HP/4-HB) pathway and AOB using the less efficient Calvin-Benson-Bassham (CBB) cycle^15^. The efficiency of these cycles is not accounted for in the TEEM model leading to an underestimation of carbon fixation for AOA. This is also reflected in the carbon balances that show a rate of ∼0.1 mol C/mol N in more balanced conditions (C and G) (Figure EV3). Although the growth curves of F and B were affected by low amounts of carbon, cells were still able to fix a comparable amount of carbon per mole of ammonia oxidized. The most limited condition, E, exhibiting the most extreme limitation is exclusively dependent on the slow release of carbon dioxide from pyruvate (Figure 1B).

This slow release of carbon induces a shift of the proteome from informational processes towards building block synthesis (Figure 3B) and therefore an increase in the relative abundance of proteins essential for the core carbon metabolism to make these building blocks, i.e. a shift from replication and active population growth towards what could be interpreted as a maintenance growth mode. This can also be seen by the high relative abundance of amino acids under carbon limitation (Figure 5).

### Responses to carbon limitation indicate key enzymes and pathways in the central carbon metabolism

Previous work in AOA^11,15^, as well as isotopic studies in *Metallosphaera sedula*^29^, an archaeon utilizing the same carbon fixation pathway, has pointed toward succinyl-CoA as the connection point between the carbon fixation and the core central metabolism via the TCA cycle, and a subsequent connection to gluconeogenesis with PckA. The results shown here strongly support this flow of carbon in the cell. Most of the proteins facilitating this flow of carbon were found to increase in relative abundance under extreme limitation. Four notable exceptions occur at PhaB (carbon fixation pathway), fumurate hydratase (FumC, TCA cycle), malate dehydrogenase (Mdh, TCA cycle), and phosphoenolcarboxykinase (PckA, gluconeogenesis) (Figure 4). In the case of PhaB, the protein did not exhibit a significant change in relative abundance between the two extreme conditions of this experiment (A and E, Figure EV10), indicating a tight regulation to keep it stably present within the cell. PhaB also represents a unique enzyme within the carbon fixation cycle as the only protein reducing, rather than oxidizing, an electron carrier and interacting with NAD^+^, while most other energetic steps rely on NADPH^15^. Differential response to the redox status of cells is often dictated by interactions with either NADH or NADPH^30^, although this balance in lithotrophs is ambiguous as NADH is not the primary input to the ETC as it is in organotrophs. The interaction of PhaB with NADH rather than NADPH, and its dissimilar response to most other carbon fixation proteins suggests that it is regulated by a stimulus other than carbon supply.

Neither PckA or FumC showed a statistically significant change in relative abundance across any carbon conditions, while Mdh, between FumC and PckA in the pathway, slightly decreased under carbon limitation (Figure EV10). Mdh converts malate to oxaloacetate and while Mdh is seen to decrease in relative abundance under carbon limitation, it was still found to be within the 10% of proteins with the highest relative abundance in all conditions, highlighting its importance. An alternative fate for malate is the conversion to pyruvate by malic enzyme (YtsJ). From a genomic perspective, this pyruvate could enter gluconeogenesis via pyruvate phosphate dikinase (PpdK). Although PpdK has been detected in previous proteomic datasets of *N. viennensis*^11^, this protein was not detected in the dataset presented here leaving PckA as the primary connector between the TCA cycle and gluconeogenesis.

Similar to Mdh, several other proteins within the carbon metabolism were found to be some of the most highly abundant proteins regardless of carbon limitation (top 10% across all conditions, Figure 5, outlined in gold). All of these above proteins represent crucial steps or connection points between core metabolic pathways that are likely highly regulated to maintain the flow of carbon within the cell. The decrease in relative abundance of Mdh suggests that it is under a different regulatory control than many other highly abundant proteins in the central carbon metabolism that were seen to increase under carbon limitation (Figure 4).

### Responses of PII regulatory proteins to carbon limitation underscore roles in central metabolic regulation

One possible regulatory mechanism that connects different metabolic pathways in the cell could come from the action of nitrogen regulatory P-II proteins. *N. viennensis* encodes for six P-II proteins in its genome^11^, which are predicted to tightly control the flow of carbon and nitrogen in the cell. Five out of six were detected in the proteomes. Two of these proteins (CnrC and CnrD), both of which increased under carbon limitation, are among the proteins with the highest relative abundance across all conditions (Figure EV11). Their high abundance, coupled with an increase under carbon limitation, suggests that their regulatory functions help to increase the flow of carbon into the cell. Conversely, CnrB was decreased under carbon limitation, and GlnB almost completely disappeared from the detected proteome (Figure EV11). A decrease in GlnB is unsurprising as its activity is known to inhibit the activity of acetyl-CoA carboxylase (Acc) in *Escherichia coli*^31^. However, this also represents a unique response in *N. viennensis*, and perhaps other archaeal autotrophs when presented with carbon limitation. In bacteria and eukaryotes, a decrease in carbon metabolites initiates an increase in GlnB to prevent the storage of carbon in fatty acids by the activity of Acc^31–33^. The opposite effect is observed here, as Acc is the key enzyme for fixing carbon in AOA rather than for storing excess carbon in fatty acids (lipids not commonly found in archaeal biomass). As such, the regulation of GlnB must be under a different mechanism than that found in other domains of life that rely on Acc for fatty acid synthesis. It is plausible that AOA rely on GlnB as a way to regulate Acc under carbon excess, but a unique regulation mechanism would still be needed to reverse this under carbon limitation. Alternatively, the consistent presence of GlnB within AOA genomes could point towards other functional roles outside of Acc inhibition. The need for AOA to decrease GlnB under carbon limitation would likely impact these other roles. The increased number of PII proteins found within AOA, with presumably different ligand binding sites, may be a way to compensate for adverse effects of GlnB down-regulation.

### Translation is a tightly controlled informational process under carbon limitation

The combined analysis of proteins and metabolites revealed additional insights into the cellular response of *N. viennensis* to carbon limitation. The protein with the highest correlations to metabolites, methionine sulfoxide reductase A (MsrA), is known to correct oxidative damage of free methionine within the cell^34^. The protein with the fifth highest correlations, a translation initiation factor (eIf-2a), is also known to interact with methionine to start translation of an mRNA sequence^35^. MsrA also correlated with inosine-5’-monophosphate dehydrogenase (GuaB), the rate limiting step of GMP synthesis^36^, the precursor for GTP and the energetic driver of the translational process^37^. A protein with a putative annotation of the same step is also one of the most abundant proteins across all conditions (NVIE_023460). GMP synthesis appears to be uniquely targeted compared to other nucleotides as most of the purine synthesis pathway, but not pyrimidine synthesis, is found within the most limited cluster (Figure 4, Figure EV12, Dataset EV1) and the rate limiting step of AMP synthesis, PurA, did not exhibit a statistically significant change between different conditions. The correlations of MsrA with eIF-2A, and GuaB, correspond strongly with the increase in relative abundance of amino acids of the metabolome and the enrichment of amino acid metabolism of the proteome under carbon limitation. Taken together this suggests translation as a bottleneck under carbon limitation and highlights key regulatory points fundamental in this process (Figure 6). The cell could also be viewed as primed for translation as almost all translation initiation factors are up-regulated under carbon limitation, while translation elongation factors and ribosomal proteins tend to be down-regulated. This might reflect the need of the cell to slow down its growth, cell division, and informational processes in response to its carbon limited environment while maintaining the molecular machinery and substrates needed to engage in translation of crucial proteins. This approach aligns with recent a metatranscriptomic study in which microbial populations responded to warming temperatures by downregulating ribosomal genes while up-regulating amino acid synthesis genes^38^. This seemingly counterintuitive approach is speculated to be a resource allocation strategy for cells under stress.

**Figure 6:**
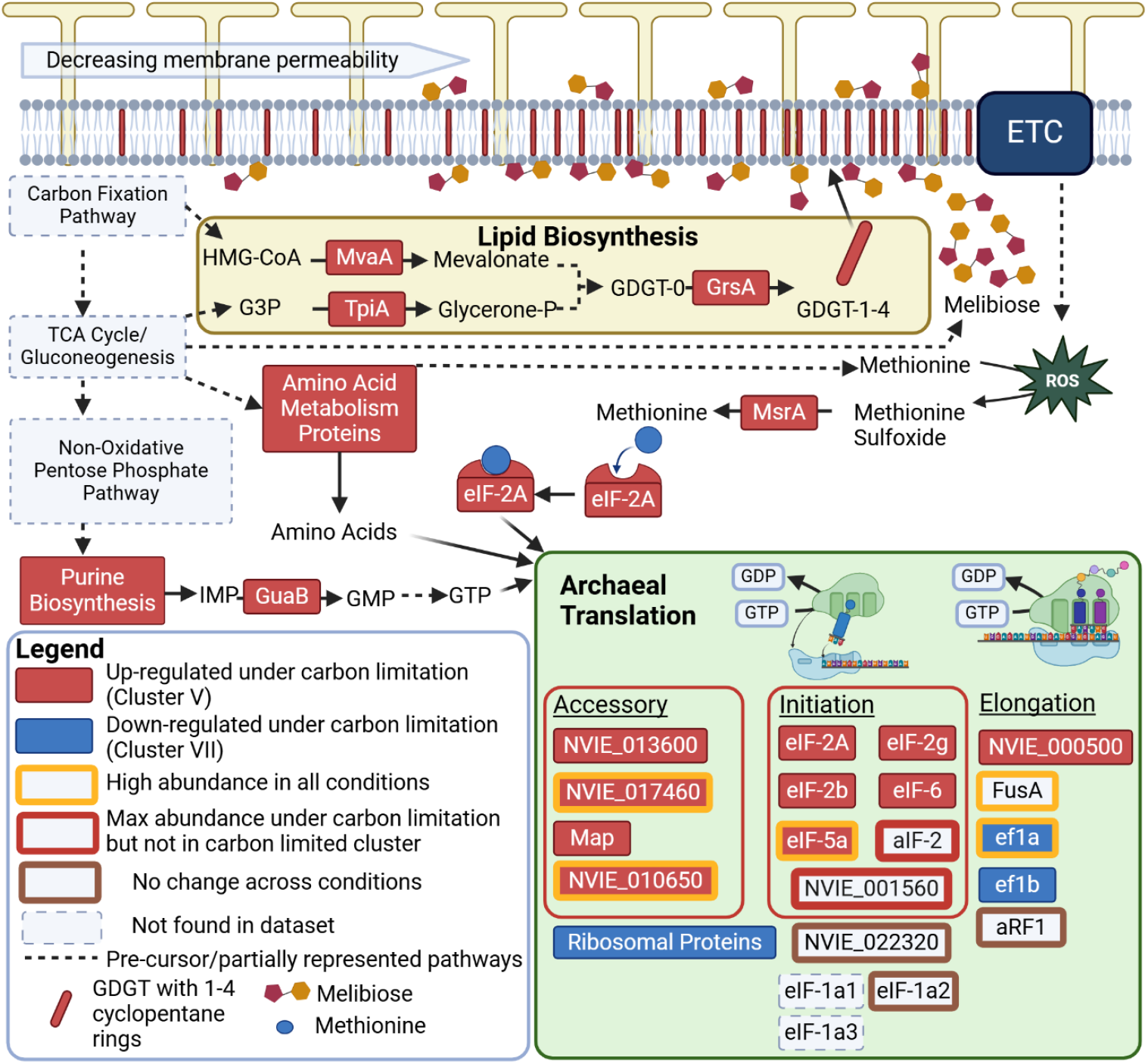
*Nitrososphaera viennensis* targeted metabolic reactions under carbon limitation. Protein abbreviations and identifiers are found in rectangular boxes. Up-regulated protein under carbon limitation is defined as proteins found within the carbon limited cluster (Cluster V) that show a statistical difference between condition E and condition A. Down-regulated proteins under carbon limitation are defined as proteins found in the carbon replete cluster (Cluster VII) that show a statistical difference between condition E and condition A. Corresponding locus tags can be found in Dataset EV1. Abbreviations: ETC, electron transport chain; HMG-CoA, 3-hydroxy-3-methylglutaryl coenzyme A; G3P, glyceraldehyde-3 phosphate; glycerone-p, glycerone-phosphate; GDGT-0, glycerol dialkyl glycerol tetraether with no cyclopentane rings; GDGT-1-4, glycerol dialkyl glycerol tetraether with 1-4 cyclopentane rings respectively; IMP, inosine monophosphate; GDP, guanosine diphosphate; GTP, guanosine triphosphate. Locus tags and accession number for proteins can be found in Dataset EV1. Created in BioRender. Hodgskiss, L. (2025) https://BioRender.com/y37o701.

A focus on translation is also a plausible explanation for the high responsiveness of NVIE_010650 from the PLS-DA analysis (Figure EV5). A structure-based search revealed that NVIE_010650 contains a PRC-barrel domain. These domains can have implications in both archaeal FtsZ cell division^39^ and ribosome maturation^40^. While *N. viennensis* encodes an FtsZ homolog, this was not detected in any of our datasets. The presence of Cdv proteins that correspond with the proteasome support previous data from *Nitrosopumilus maritimus* that AOA do not divide with an FtsZ-like system^27^, thus ruling out a role of NVIE_010650 in cell division. Although significant differences can be seen in the amino acid alignments (Figure EV13), the structural search found the closest protein candidates to likely be involved in ribosome maturation (Dataset EV1).

### Melibiose as a lipid membrane stabilizer

A stark response of *N. viennensis* to carbon limitation was the presence of melibiose, a disaccharide which was not detected during growth in carbon-replete conditions. The pattern of melibiose is also distinct from other sugars (Figure EV7). Sugar metabolism within AOA is not well studied, but the investment of producing a disaccharide sugar under extreme carbon limitation suggests that it is an important molecule for the functioning of the cell under carbon limitation stress. One plausible role of melibiose could be as a polar head group to archaeal lipids. AOA are known to include a large amount (∼40%) of dihexose head groups^41–44^, and while the full molecular structure of these head groups is unknown, glucose and galactose, the monosaccharides that comprise melibiose, are known to be found in archaeal lipid analyses of intact polar lipids^45,46^. An increase in dihexose head groups could be beneficial for the cell by tightening the membrane^47,48^ and preventing the escape of vital carbon metabolites. However, currently only the addition of monosaccharides to lipids in archaea has been documented^49,50^ rather than pre-made dihexose molecules. Alternatively, melibiose could also be interacting with the membrane via a stabilizing effect as seen for disaccharides in other organisms^51,52^. A membrane adjustment under carbon limitation is additionally supported by an increase in the relative abundance of GrsA (NVIE_028400) one of the proteins responsible for cyclization of GDGT lipids^53^ (Figure 6). Increased cyclization of lipids in organismal membranes is known to reduce membrane fluidity and has been positively correlated with reduced growth rate in the marine AOA *Nitrosopumilus maritimus*^54,55^. AOA have been observed to release 5-15% of their fixed carbon as dissolved organic carbon metabolites during growth^28^ some of which could be escaping the cell via passive diffusion through the membrane^56,57^. Reduced membrane fluidity would decrease permeability and likely reduce the loss of crucial carbon metabolites (particularly hydrophobic amino acids^58^) to passive diffusion of metabolites through the archaeal membrane^59^.

### Choice of ROS scavenger triggers unique metabolic strategies for reactive species detoxification

Apart from carbon limitation, distinct differences in proteomes and metabolomes of *N. viennensis* were detected between growth conditions that used either pyruvate or catalase as the exogenous ROS scavenger. With pyruvate, an increase in several proteins involved in maintaining redox homeostasis was observed (Figure 7), while with catalase there was the exclusive presence of the sugars glucose, maltose, and trehalose (Figure EV7). While both catalase and pyruvate have been reported to detoxify ROS and RNS^60–62^, the distinct cellular reactions can most likely be explained by the accessible sites of the ROS scavengers. Catalase constitutes a large protein that, due to its size, is not likely able to pass through the S-layer to the pseudo-periplasm where the vast majority of ROS and RNS will be produced from the electron transport chain. This likely leads to an accumulation of ROS/RNS in the pseudo-periplasm where the critical steps of the ammonia oxidation process are likely taking place. In response to this, the production and excretion of sugars, particularly trehalose and maltose, both of which are comprised by glucose, could be exported to the pseudo-periplasm to deal with this accumulation of ROS. The production of these sugars, which is only observed in the catalase conditions, would also help to relieve ROS stress within the cell (Figure 7). This hypothesis is supported by the ROS relieving observation of sugars in other organisms^63–65^, including *N. viennensis* (albeit it to a lesser extent than pyruvate; supp. material Tourna et al. 2011^20^), and the correlation of all three sugars with an ABC efflux transporter protein that could transport the sugars to the pseudo-periplasm. The production of these sugars may also be connected with a putative TetR transcriptional regular (NVIE_002180) that was also positively correlated with all three.

**Figure 7:**
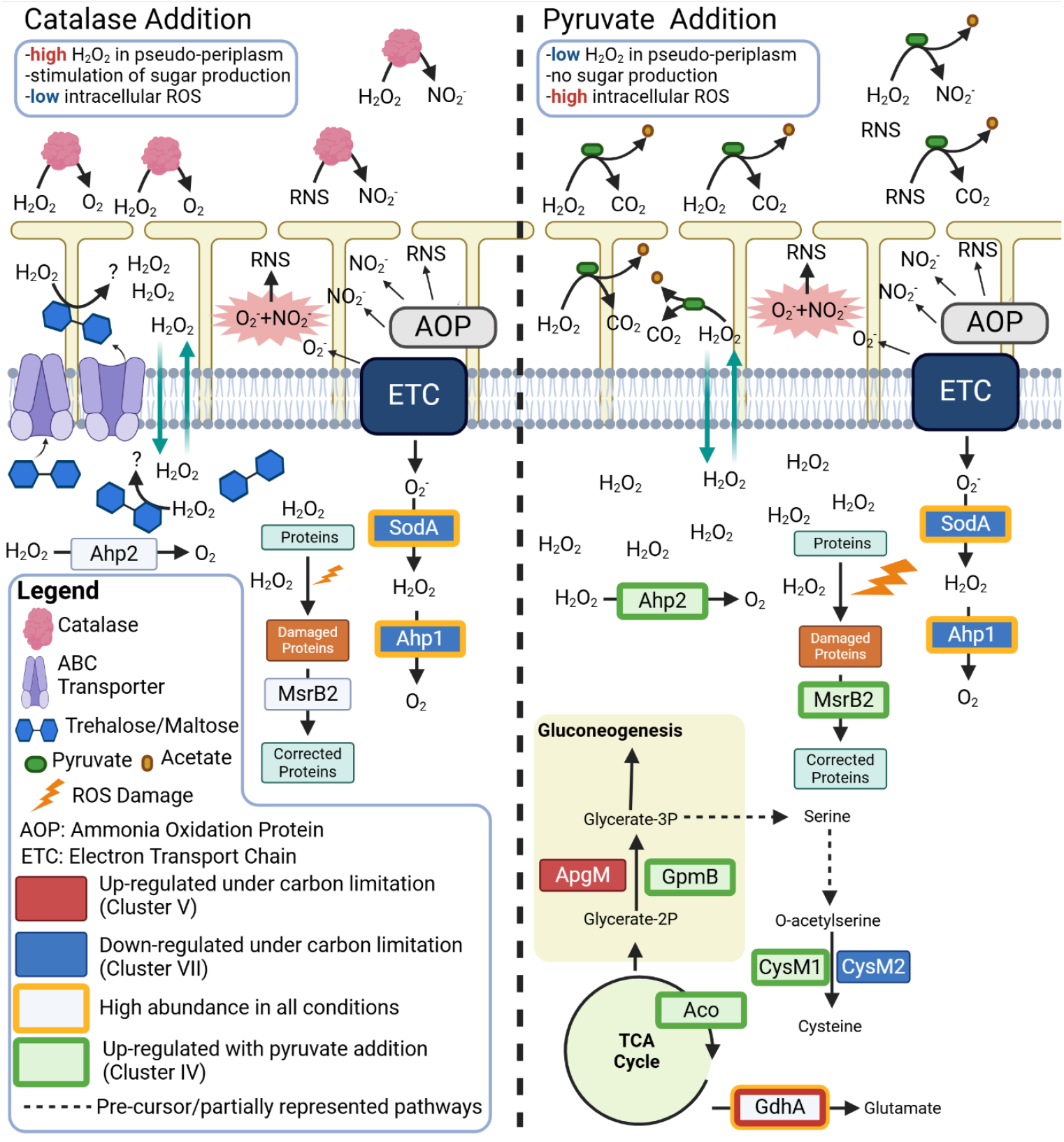
*Nitrososphaera viennensis* model of reactive oxygen species (ROS) coping strategies. Protein abbreviations and identifiers are found in rectangular boxes. Up-regulated protein under carbon limitation is defined as proteins found within the carbon limited cluster (Cluster V) that show a statistical difference between condition E and condition A. Down-regulated proteins under carbon limitation are defined as proteins found in the carbon replete cluster (Cluster VII) that show a statistical difference between condition E and condition A. Proteins upregulated with addition of pyruvate were found in Cluster (IV) and showed a statistical difference between at least two conditions. Corresponding locus tags can be found in Dataset EV1. Abbreviations: AOP, unknown ammonia oxidation protein(s); ETC, electron transport chain; N, non-reactive nitrogen species; glycerate-2P, glycerate-2 phosphate; glycerate-3P, glycerate-3 phosphate; H_2_O_2_, hydrogen peroxide; O_2_, oxygen; NO_2_^-^, nitrite; RNS, reactive nitrogen species; O_2_^-^, superoxide. Locus tags and accession number for proteins can be found in Dataset EV1. Created in BioRender. Hodgskiss, L. (2025) https://BioRender.com/x36a147.

Conversely, most of the ROS/RNS in the pseudo-periplasm of the pyruvate containing conditions is likely directly detoxified by pyruvate, a molecule small enough to fit through the S-layer lattice. The lack of ROS stress in the pseudo-periplasm likely negates the need for the production and excretion of sugars. While this solves the ROS problem in the pseudo-periplasm. The lack of sugar production within the cell leaves a greater amount of intracellular ROS when compared to conditions that have catalase. The increase in Ahp2 (NVIE_011770) to detoxify ROS/RNS and MsrB2 to correct damaged proteins is likely a response to this. The up-regulation of GpmB and one of two cysteine synthases (CysM1, NVIE_017430) may also be reinforcing the production of cysteine, a particularly susceptible amino acid to ROS and RNS stress (additional info in Supp. Material). A re-direction of carbon in the cell is also supported by the response of aconitase (Aco, NVIE_029450) (Figure7). The aconitase in AOA has previously been reported to be evolutionarily related to mitochondrial aconitase and possibly sensitive to the redox state of the cell^21^. From these observations, it is likely that SodA and Ahp1 are the primary ROS detoxifcation proteins in *N. viennensis*, while Ahp2, MsrB2, and specific carbon metabolic response are utilized when ROS stress increases intracellularly. The strategy of keeping a primary Ahp highly expressed with additional Ahp proteins up-regulated when needed is also seen in marine AOA under different ROS stresses (Bayer et al. 2019^66^).

It is tempting to speculate that the pyruvate could be driving the slight up-regulation of Aco for carbon assimilation. An early study of *N. viennensis* showed incorporation of C^13^ labeled pyruvate into biomass (5.85-9.54% of total carbon) along with a production of C^13^ labeled CO_2_ accounting for 6.46-7.09% of CO_2_ in the culture headspace^20^. It was later proven that hydrogen peroxide is detoxified by pyruvate leading to a release of CO_2_^24^. The ROS detoxifying nature of pyruvate and the lack of an endogenous catalase in many AOA, including *N. viennensis*, explains the substantial growth rate increase that is stimulated by some organic acids^20^. This new information casts doubt on a mixotrophic nature of *N. viennensis*. The previously labeled biomass was likely the result of the cell fixing C^13^ labeled CO_2_ that had been cleaved from the fully C13 labeled pyruvate as evidenced by its presence in the culture headspace. The direct incorporation of organic carbon into AOA biomass has been suggested for other clades based on genomic comparisons but has yet to be substantially confirmed^67^. For these reasons, a direct incorporation of pyruvate into biomass is yet to be shown for *N. viennensis* and is not assumed to be happening in this study.

### An archaeal chemolithoautotrophic response to carbon limitation

As a chemolithoautotroph, the response of *N. viennensis* to carbon limitation offers a unique look at this nutrient stress without directly impacting the energy source of the cell. This is not possible in well studied heterotrophic organisms where the carbon and energy source are directly intertwined. Even in the more similar energetic system of photolithotrophs, carbon limitation is usually achieved when light, and therefore energy, is not available, again directly tying carbon to the energy source of the cell. As lithoautotrophs, a decoupling of energy and carbon, and therefore the associated cellular response, is achievable in AOA. Although growth was slowed under carbon limitation, the significant reduction of nitrification observed in the cell is a direct response to carbon limitation rather than energy as ammonia and oxygen were readily available. In *N. viennensis*, the functional response to carbon limitation observed with this systems approach emphasized its strictly autotrophic lifestyle, as it consisted of up-regulating and maintaining its standard core metabolism. This up-regulation was seen even with enzymes that are already at a constitutively high level.

In addition to an observed response in core metabolism, factors involved in cellular translation, and in particular translation initiation, were associated with carbon limitation in *N. viennensis*. A regulation of translation and amino acid synthesis under carbon limitation, or nutrient stress, has been observed in species across all three domains of life. In eukaryotes this is achieved through the highly regulated AMPK/TOR signaling system^68–70^, while bacteria rely primarily on the stringent response and the regulator ppGpp^71,72^. Archaea do not have an AMPK/TOR signaling cascade, and while *N. viennensis* contains an up-regulated putative SpoT/RelA gene that could initiate ppGpp synthesis, a stringent response mediated by ppGpp has not been observed in archaea. While some archaea appear to have a stringent response, it occurs without the production of ppGpp and without a drop in GTP levels^73,74^, making them unique to most bacterial stringent response systems. Although translational responses to ppGpp have since been documented in the archaeon *Sulfolobus* (now *Saccharolobus*) *solfataricus*^75^, the effectiveness of ppGpp seemed to be diminished and more specialized. Regardless, specific responses in prokaryotes to carbon limitation range from breaking down existing proteins for amino acids (*S. acidocaldarius*)^76^, maintaining a ribosomal pool (*E. coli*)^77^, and ensuring translational fidelity (*T. kodakarensis*)^78^. All of these observed responses ensure the ability of translation to proceed in the cell. In this study, *N. viennensis* was found to maintain translational activity through a concerted effort of de novo synthesis of amino acids and GTP, decreasing membrane permeability, and simultaneously up-regulating translation initiation factors.

In summary, the response of *N. viennensis* to carbon limitation illustrates a tight control of translational processes as seen in other organisms across the tree of life. However, the exact regulatory mechanisms of this response remain elusive as the typical nutrient limitation response mechanisms in eukaryotes and bacteria are either not present or not strongly observed within the archaeal domain, likely reflective of a domain with metabolic pathways similar to bacteria but informational machinery more akin to eukaryotes. Alongside translational and core metabolism regulation, *N. viennensis* demonstrated the ability to respond to environmental stimuli in a precise manner by modifying its metabolome and/or proteome. The modelling approach combined with proteomics and metabolomics used here allowed for the detection of fine-tuned responses in *N. viennensis* that help explain its success in a wide range of soil habitats.

## Methods

Detailed methods on culturing of *Nitrososphaera viennensis*, chemical analysis (nitrite/ammonium/inorganic carbon measurements), extraction protocols (combined protein/metabolite extraction with methanol/chloroform/water), and data analysis can be found in Supplementary Material.

### Thermodynamic Model

A thermodynamic model for the metabolism of *N. viennensis* was constructed based off of the Thermodynamic Electron Equivalents Model 2 (TEEM2) method^22^.

The electron donor reaction was calculated as the oxidation of ammonia to nitrite. Ammonia was chosen for the half reaction instead of ammonium as it is assumed that ammonia, and not ammonium, is the true substrate for the AMO complex^79^ that begins the ammonia oxidation pathway:

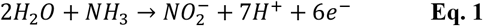

The reaction was then normalized to one electron and arranged in the reducing direction:

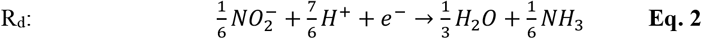

Oxygen reduction was used as the half reaction for the electron acceptor:

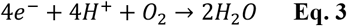

This equation was also normalized to one electron:

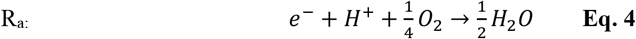

Carbon and nitrogen measurements of washed *N. viennensis* determined the C:N molar ratio to be 3.85:1. Although molar ratios of oxygen and hydrogen were not determined, a biomass composition of C_3.85_H_6.69_O_1.78_N was used. This was based off of values for *Escherichia coli* that had the same C:N molar ratio of 3.85:1^23^ (pg. 129). The biomass composition was used with the following formula to estimate a half reaction for biomass formation:

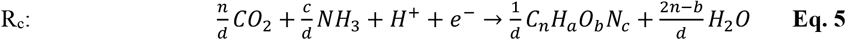

where *d* = 4*n* + *a* − 2*b* − 3*c* .

This equation is modified from Rittmann and McCarty^23^ (pg. 137). Typically, a generic biomass concentration would include NH_4_^+^ rather than NH_3_ to represent a nitrogen source. As the nitrogen source and electron donor in AOA are the same, the equation was modified to use NH_3_ based on the physiological observation that the vast majority of nitrogen consumed by *N. viennensis* is used for nitrite production and would therefore be utilized in the form of ammonia. The original equation also includes HCO_3_^-^ to balance the positive charge of ammonium. Therefore, CO_2_ was used in place of HCO_3_^-^ as a counter-charge balance was not needed. Although the actual substrates for AOA biomass are NH_4_^+^ and HCO_3_^-^, the use of NH_3_ and CO_2_ are more consistent when taking into account the electron donor (NH_3_) from Eq. 1 and 2 and do not alter the biomass equation as each contains the same amount of nitrogen or carbon respectively. Utilizing the estimated biomass composition and the modified equation gives the following:

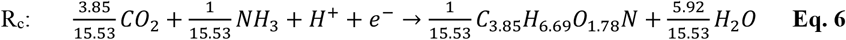

Once the necessary reactions were obtained, ΔG’ values were calcualted. Gibbs free energy was calculated for a temperature of 25°C and pH of 7.0. These are not the exact temperature of the growth conditions for *N. viennensis* (42°C and pH ∼7.2 respectively). However, standard conditions were used as these are the parameters used to calculate the Gibbs free energy for the reduction of oxygen.

Oxygen reduction: 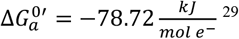

As this is an empirical value based on a hydrogen potential with platinum, it is not readily adjustable. For consistency, the same parameters were used for the calculation of Gibbs free energy for the half reaction of ammonia to nitrite. Gibbs free energy of formation was used for the participating chemical species:

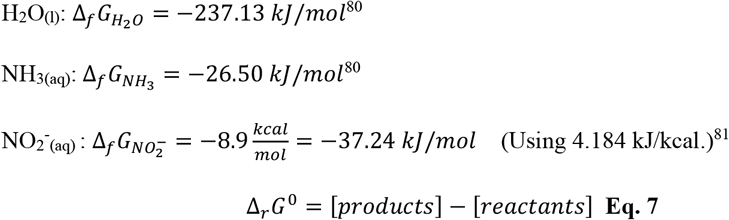

Using Equation 2:

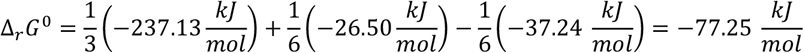

To adjust for pH of 7.0:

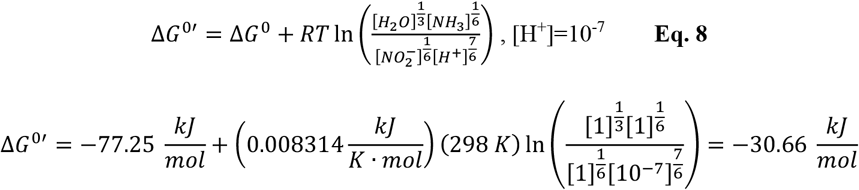

Ammonia oxidation in reducing direction: 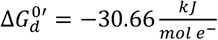

For calculation of model:

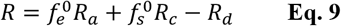

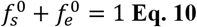

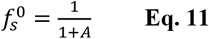

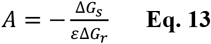

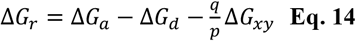

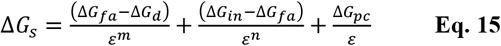

where ΔG_a_ is the Gibbs free energy for the electron acceptor half reaction, ΔG_d_ is the Gibbs free energy for the electron donor half reaction, q represents the number of monooxygenase reactions per substrate, p is the number of electron equivalents per mole of substrate, ΔG_xy_ is the reduction potential for NADH oxidation (representative of the energy input for the monooxygenase reaction; -219.2 kJ/mol^22^), ΔG_fa_ is the reduction potential for a formaldehyde half reaction, ΔG_in_ is the Gibbs free energy for the reduction potential of acetyl-CoA (representative of carbon activation for autotrophs; 30.9 kJ/mol^22^) ΔG_pc_ is the Gibbs free energy for intermediate conversion to cells, ε is the energy transfer efficiency, m is 1 if ΔG_fa_ is greater than 0 and equal to n otherwise, and n is 1 if ( ΔG_in -_ ΔG_d_)>0 and m=n^22^.

In the case of *N. viennensis*, ΔG_fa_ =0 as no formaldehyde is involved and ( ΔG_in -_ ΔG_d_)>0. This causes m=n=1. The Equation 13 for ΔG_s_ therefore simplifies to:

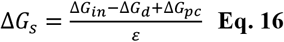

The value of p is equal to 6 (6 electrons per ammonia, Eq. 1), q is 1 (one monooxygenase reaction per ammonia), and ΔG_pc_ is the energy to synthesize cells:

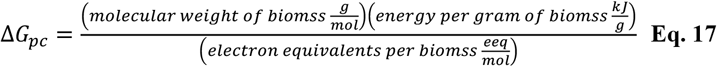

Molecular weight of C_3.85_H_6.69_O_1.78_N = 95.37 g/mol

Energy per gram of biomass=3.33 kJ/g ^23,82^

Electron equivalents per biomass= 15.53 eeq/mol (Equation 6)

Input values for Equation 17:

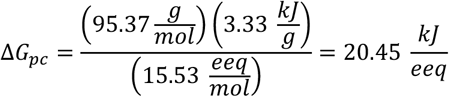

The energy transfer efficiency, ε, is estimated to be 0.57 as observed for other autotrophs^22^.

Input values for Equation 16:

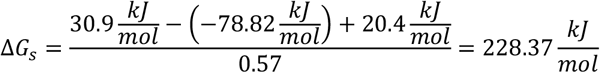

Input values for Equation 14:

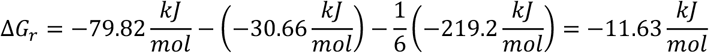

Input values for Equation 13:

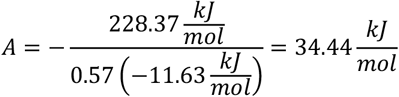

Input values for Equation 11:

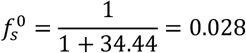

Input values for Equation 10:

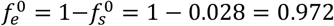

Input values and equations (R_a_ as Eq. 4; R_d_ as Eq. 2; R_c_ as Eq. 6) for overall reaction (Equation 9):

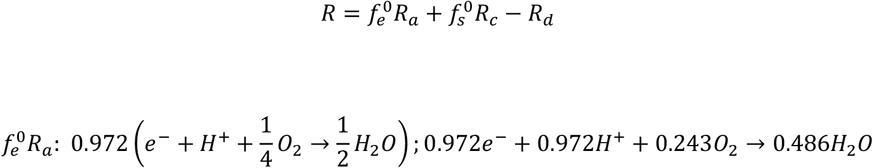

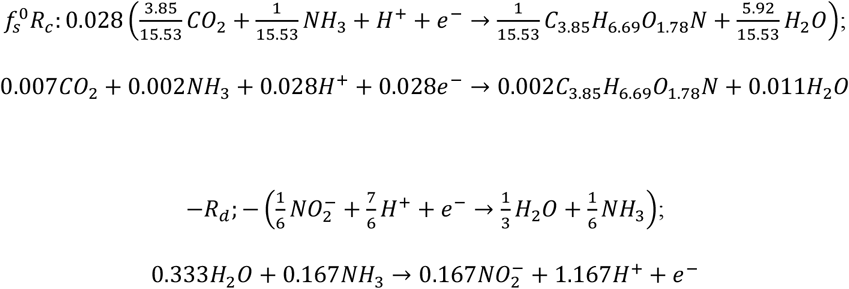

Combine all equations and normalize to 1 mol of ammonia:

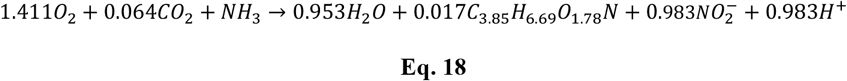

Equation 18 normalized to 2 mol ammonia:

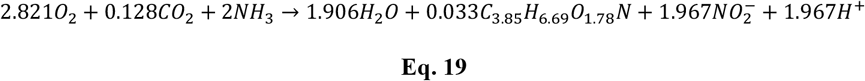

### Experimental Setup for Carbon Conditions

With a nitrogen input concentration of 2 mM (ammonia/ammonium), a theoretical carbon threshold of 0.128 mM was used for the experimental design of growth curves with varying levels of inorganic carbon: 2 mM carbon with catalase (A); 0.75 mM carbon with catalase (D) 0.25 mM carbon with pyruvate (G); 0.25 mM carbon with catalase (C); 0.1 mM carbon with pyruvate (F); 0.1 mM carbon with catalase (B); and 0 mM carbon with pyruvate (E). This spread of carbon concentrations produced a gradient from high to low amounts of available inorganic carbon (Figure 1A). The reaction of pyruvate with ROS would create additional available carbon for the cell. In the case of the 0 mM inorganic carbon concentration, this created a slow release of carbon substrate over time. The pyruvate conditions with 0.25 mM and 0.1 mM of added carbon would therefore have slightly more available carbon than their catalase counterparts. The combination of all conditions gives a gradient of carbon concentrations starting from 2 mM and dropping down to just above 0 mM.

## Data and Code Availability

The mass spectrometry proteomics data have been deposited to the ProteomeXchange Consortium via the PRIDE^83^ partner repository with the dataset identifier PXD060602 and 10.6019/PXD060602. Metabolomic data has been deposited to the MetaboLights database under accession number MTBLS11689.

## Acknowledgements

We thank Lena Fragner for technical guidance during extraction of metabolites and proteins, Sonja Tischler for mass spectrometry technical support, and Hubert Kraill for assistance with running the elemental analyzer. We also thank Lisa Fürtauer, Maria Pacheco, Jakob Weiszmann, and Martin Brenner for very valuable and helpful discussions during the data analysis. We are also appreciative of Dr. Thomas Rattei and team of the Division of Computational Systems Biology (CUBE) for providing maintenance and access to the Life Science Computer Cluster (LiSC) at the University of Vienna.

