## Supplementary material for "Metabolic Response of a Chemolithoautotrophic Archaeon to Carbon Limitation": Dataset EV1

**Contents:**

Supplementary Discussion

Supplementary Methods

Supplementary Figures S1-S12

Supplementary Data (Datset_S1, included Excel sheet)

### Supplementary Discussion

*Proteins highly abundant in all conditions*

Enrichment analysis according to archaeal clusters of orthologous groups (arCOGS)[1] of these highly abundant shared proteins showed a high number of proteins involved in: energy production and conversion (arCOG C) and translation, ribosomal structure and biogenesis (arCOG J) (Figure EV14). Proteins implicated in energy production included AmoB (ammonia monooxygenase enzyme subunit B), NirK, and subunits of ATP synthase (AtpA, AtpC, and AtpE). Numerous proteins involved in carbon metabolism were detected including aspartate semialdehyde dehydrogenase (Asd; carbon fixation cycle), glutamate dehydrogenase (GdhA; ammonium assimilation), malate dehydrogenase (Mdh; TCA cycle) and bifunctional fructose-1,6-bisphosphate/aldolase (Fbp; gluconeogenesis). Although these proteins represent the most highly abundant proteins in all conditions, the majority of them (89%) were also seen to change in relative abundance across conditions suggesting that they are still under regulatory control with respect to carbon concentration.

*Proteins with no statistical change across conditions*

Proteins that did not change in response to carbon concentrations included ribosomal proteins and selected proteins involved in purine and pyrimidine metabolism but were not enriched for any arCOG category (Figure EV14). AmoB, AmoY (a newly identified subunit of the archaeal ammonia monooxygenase[2]), and NirK were also included in this category underscoring their importance to cellular function regardless of carbon limitation. Other proteins involved in purine metabolism (PurU, PurA, AdkA, PurD, GuaA ), pyrimidine metabolism (ThyX, PyrE, PyrF. PyrB), the non-oxidative pentose phosphate pathway (RpiA, Tal), and the conserved connection point between C-3 and C-4 metabolism[3] in AOA (PckA, ATP-dependent phosphoenolpyruvate carboxykinase) were also found to remain constant across the different growth conditions. Many of these are involved in carbon metabolism and the production of vital metabolites, i.e. nucleotides.

*Core metabolism response to pyruvate addition*

A response at specific points within the core metabolism was also observed that could be explained by the presence of additional redox stress from ROS or RNS (Figures 4 and 7). Within gluconeogenesis, a redundancy in the conversion of glycerate-2P to glyerate-3P exists in *N. viennensis* with the proteins ApgM and GpmB both facilitating this step. While ApgM is higher in all conditions and responds to carbon limitation, the additional up-regulation of

GpmB specifically under pyruvate would also direct more carbon to glycerate-3P, the precursor for serine and therefore cysteine, an amino acid that is commonly impacted by ROS/RNS stress. In support of this, a redundant putative cysteine synthase (CysM1) was also found within this cluster, suggesting a response of the cell to replace commonly damaged amino acids (i.e., cysteine). A slight up-regulation of aconitase (Aco) within the TCA cycle is also seen in pyruvate conditions which may also support the flow of carbon towards alpha-ketoglutarate and the assimilation of ammonium, a necessary step for amino acid production to replace damaged proteins and amino acids, including cysteine. The response of aconitase is supportive of the acquisition of a mitochondrial-type aconitase (Aco) in AOA[4] that is likely responsive to the redox status of the cell[4,5] (Figures 4 and 7).

### Supplementary Methods

*Carbon and Nitrogen Content of Cell*

Cells were grown under standard conditions with pyruvate and harvested in late exponential growth phase by centrifugation (16,000 x*g* at 4°C for 1 h). Cell pellets were washed twice with FWM (no additions of bicarbonate or ammonium), dried for 1h (vacuum concentrator) and stored in a desiccator until further processing. Subsequently, cell pellets were weighed and packed into tin capsules prior to being analyzed on a CHNS elemental analyzer (Vario MICRO cube, Elementar). The instrument was calibrated with acetanilide following manufacturer protocols.

*Culture Conditions*

Cultures of *N. viennensis* were grown in closed 1 liter pressure safe Schott bottles closed with a grey bromobutyl rubber stopper (GL45 Duran) with 580 mL of fresh water medium (1g/L NaCl, 0.4 g/L $MgCl_2 \cdot 6H_2O$, 0.1 g/L $CaCl_2 \cdot 2H_2O$, 0.2 g/L $KH_2PO_4$, 0.5 g/L KCl). The choice of rubber stopper is critical as *N. viennensis* will not grow when certain other rubber stoppers are used. The following was added to FWM: 600 µL trace element solution (100 mM HCl (~12.5M), 0.5 mM $H_3BO_3$, 0.5 mM $MnCl_2 \cdot 4H_2O$, 0.8 mM $CoCl_2 \cdot H_2O$, 0.1 mM $NiCl_2 \cdot 6H_2O$, 0.01 mM $CuCl_2 \cdot H_2O$, 0.5 mM $ZnSO_4 \cdot 7H_2O$, 0.15 mM $Na_2MoO_4 \cdot 2H_2O$; autoclaved, stored in dark), 600 µL vitamin solution (0.02 g/L biotin, 0.02 g/L folic acid, 0.1 g/L pyridoxine HCl, 0.05 g/L thiamine HCl, 0.05 g/L riboflavin, 0.05 g/L nicotinic acid, 0.05 g/L DL pantothenic acid, 0.05 g/L P aminobenzoic acid, 2 g/L choline chloride, 0.01 g/L vitamin B12; adjusted pH to 7 with KOH, filter sterilized, stored in dark), 600 µL 7.5 mM FeNa-EDTA (pH 7), and 6 mL HEPES buffer solution (24 g/L NaOH, 119.2 g/L HEPES; pH ~7.6, filter sterilized).

For all cultures, kanamycin was added to a final concentration of 100 mg/mL and $NH_4Cl$ was added to a final concentration of 2 mM.

A catalase stock solution was made by dissolving catalase from bovine liver (Sigma Aldrich, powder, 2000-5000 units/mg protein (average of 3500unit/mg)) in a potassium phosphate buffer solution (0.6528 g/L $KH_2PO_4$, 7.83 g/L $K_2HPO_4$, pH adjusted to 7) at 1 mg/mL. Catalase was filter sterilized before use and added at 50 U/mL final concentration[6]. Cultures grown with catalase were observed to have some translucent flocs due to the addition of catalase. These flocs formed upon addition of catalase and were therefore not attributed to be a product of cell growth. Cultures using pyruvate as a ROS scavenger had pyruvate added to a final concentration of 0.3 mM.

Inorganic carbon was added from freshly made solutions of filter sterilized 0.5M sodium bicarbonate. Based on preliminary tests amounts of added bicarbonate are overestimated (i.e. a culture with assumed 2 mM of bicarbonate is actually 1.67 mM bicarbonate). This discrepancy is likely due to the exchange of carbon dioxide between the gas and aqueous phases. The actual concentration of a freshly prepared 0.5 M stock solution was determined to be 0.415 mM. This concentration was used to calculate the amount of stock solution to add to cultures. Therefore, for 600 mL cultures, the following amounts of freshly made 0.5M sodium bicarbonate solution was added: 2 mM carbon culture was supplied with 2.87 mL, a 0.75 mM culture was supplied with 1.08 mL, a 0.25 mM culture was supplied with 0.36 mL, and a 0.1 mM culture was supplied with 0.14 mL.

All cultures were inoculated with 1.5 mL (0.25% inoculum) of an actively growing culture in late exponential phase. Final volume after additions and inoculation comes to approximately 600 mL.

After inoculation, the headspace of cultures was flushed with an artificial gas mixture of 80% nitrogen 20% oxygen to remove any residual carbon dioxide in the head space. Immediately after flushing the head space, a water sample was taken from the aqueous phase to determine the total starting carbon concentration (see below). Cultures were then grown in a 42°C incubator on a shaker set at 80 rpm.

Cultures were tracked by taking periodic nitrite measurements. Cultures were harvested when late exponential phase was reached but before entering stationary phase. At the end of a growth curve, samples were taken for gas analysis and inorganic carbon analysis of liquid. One milliliter was collected and spun down for 1 hour at 16,000xg and 4°C. The supernatant was removed and the pellet was stored at -70°C until DNA could be extracted. The rest of the culture was spun down in 250-300 mL volumes using a large Sorvall centrifuge for 45 minutes at 16,100xg at 4°C. Supernatant was poured off and the pellet was resuspended in remaining media and transferred to a new 1.5 mL Eppendorf tube. Cells were then concentrated by centrifuging at 16,100xg for 30 minutes and 4°C. As much supernatant as possible was removed without disturbing the cell pellet. Eppendorf tubes were weighed before and after cells were added to obtain an approximated wet biomass weight. Pellets were then frozen at -70°C until further metabolite and protein extraction.

*Ammonium Measurements*

Ammonium was measured colorimetrically. 200 µL of sample was mixed with 400 µL of fresh water medium (FWM) in an Eppendorf tube. 300 µL of color reagent ( 5.18 mM sodium salicylate, $2.15 \times 10^{-5}$ M sodium nitroprusside, 0.1 M NaOH; freshly prepared) was added followed by 120 µL of oxidation solution ( $3.01 \times 10^{-5}$ M dichloroisocyanuric acid; freshly prepared). Samples were shaken and incubated in the dark for 30 minutes. After 30 minutes, 200 µL of sample were pipetted into wells in a micro titer plate and absorbance was measured at 660 nm using a Tecan-Sunrise plate reader. Standard curves were made using a stock solution of 1 M $NH_4Cl$.

*Nitrite Measurements*

Nitrite was measured colorimetrically. 10 or 20 µL of sample was mixed with 790 or 780 µL fresh water medium (FWM) respectively and 200 µL of a sulfanilamide/NED reagent (150 mL ortho-phosphoric acid, 10 g sulfanilamide, 0.05 g α-naphtylethylendiamindihydrochloride in 1 L water; stored at 4°C in the dark) in a 1 mL Eppendorf tube. Samples were shaken and then stored in the dark for at least 10 minutes. After 10 minutes, 200 µL of each sample was pipetted into wells of a micro titer plate and absorbance was measured at 545 mM using a Tecan-Sunrise plate reader. Standard curves were made from a stock solution of 1 mM $NaNO_2$ (i.e. 1 mM $NO_2^-$ : 20 µL stock + 780 µL FWM; 0.8 mM $NO_2^-$ : 16 µL stock + 784 µL; 0.6 mM $NO_2^-$ : 12 µL stock + 788 µL; etc.).

*Dissolved Inorganic Carbon Analysis*

10 mL of sample were taken at the beginning (sterile, with flame and needle) and end of each culture to determine starting and ending dissolved inorganic carbon (DIC) concentration in the aqueous phase. Samples were filtered to remove cells and stored at -70°C until the inorganic carbon could be measured. DIC data were measured using a Shimadzu TOC -LCPH analyzer equipped with an DIC reaction vessel containing a reaction solution of phosphoric acid of about 25% (weight%) and an ASI-L autosampler. The samples were injected into the reaction vessel where all inorganic carbon is converted to carbon dioxide which in the following is volatized by the sparging process (synthetic air, carbon dioxide free gas) and detected by a NDIR detector. Each sample is measured three times with an injection volume of 100 micro-liter each. The final results are the corresponding mean values of the three injections per sample.

*Gas Chromatography Analysis*

At the time of harvest, head space pressure was measured and 40-50 mL of head space gas was removed using a 50 mL syringe and transferred to a glass 120 mL serum bottle pre-filled with $CH_4$ at 1 atm. Mixed gas samples were then analyzed using an Agilent Gas Chromatograph (Agilent 7890A GC, Agilent Technologies, Santa Clara, CA, USA) using a thermal conductivity detector. Gases were separated at 170°C using helium as the carrier gas. The reference flow setting was 10 mL/min. The makeup flow was set to 1 mL/min.

The following gases were used: $H_2$, $CO_2$, $N_2$, $CH_4$, and $H_2N_2CO_2$ (mix ratio 7:1:1). All gases were from Air Liquide GmbH, Schwechat, Austria. The standard test gas used for gas chromatography (GC) comprised the following composition: 0.01% volume $CH_4$ and 0.08% volume $CO_2$ in $N_2$ (Messer GmbH, Wien, Austria).

*Carbon Balance*

The total amount of carbon consumed by each culture can be approximated by the equation:

$$C_{con} = C_{avail} - C_{aq} - C_g \qquad \textbf{Eq. 20}$$

where $C_{con}$ is total carbon consumed, $C_{avail}$ is amount of available inorganic carbon, $C_{aq}$ is the aqueous inorganic carbon concentration at the end of the experiment, and $C_g$ is the gaseous inorganic carbon concentration at the end of the experiment. $C_{avail}$ can be calculated by:

$$C_{avail} = C_{aq,0} + C_{g,0} + C_{pyr} \quad \textbf{Eq. 21}$$

where $C_{aq,0}$ is the initial aqueous inorganic carbon concentration, $C_{g,0}$ is the initial gaseous inorganic carbon concentration, and $C_{pyr}$ is the amount of inorganic carbon contributed from the interaction of pyruvate and ROS. In all cultures, the $C_{g,0}$ is 0 as the head space of the cultures is flushed while $C_{aq,0}$ is represented by the initial IC measurement taken. In catalase cultures, $C_{pyr}$ is 0, while in pyruvate cultures, $C_{pyr}$ is estimated by:

$$C_{pyr} = [NO_2^-] * \frac{0.005\ mMH_2O_2}{0.5\ mM\ NH_4^+} * \frac{1\ mM\ NH_4^+}{0.983\ mMNO_2^-} * \frac{1\ mM\ CO_2}{1\ mM\ H_2O_2} \quad \textbf{Eq. 22}$$

where $[NO_2^-]$ represents the mM of nitrite produced by the culture. The ratio of 0.0045 mM $H_2O_2$/ 0.5 mM $NH_4^+$, is based off estimates of hydrogen peroxide production of *N. viennensis* from supplementary material in Kim et al. (2016) [6], 1mM $NH_4^+$/0.983 mM $NO_2^-$ is based off of the growth equation calculated in this manuscript, and 1 mM $CO_2$/1 mM $H_2O_2$ is based off the assumption of stoichiometric release of carbon dioxide from pyruvate as it interacts with hydrogen peroxide.

*DNA Extraction*

DNA was extracted from 1 mL of each culture taken at the time of harvest using the NucleoSpin Soil kit from Machery-Nagel and by following instructions for genomic DNA extraction from soil using lysis buffer SL1. Extracted DNA was eluted into 60 µL of elution buffer SE (5 mM Tris/HCl, pH 8.5). Concentrations were measured using a Qubit DNA Assay with 5 µL (high sensitivity protocol).

*Combined Protein and Metabolite Extraction*

Methods of a combined protein and metabolite extraction were followed based on those of Ott et al. (2019) [7]. Cell pellets were thawed on ice and resuspended in 500 µL methanol:chloroform:water (MCW, 2.5:1:0.5). Water for all extractions came from a MilliporeSigma Milli-Q Reference A+ System (MilliQ water). Cells and undissolved pellet were transferred to a Lysing Matrix B tube with Lysing Matrix B (MP Biomedical) filled to 2-3 mm above the cone shape at the bottom of the tube. As an internal control and extraction standard, pentaerythritol (PE) and phenyl-β-glucopyranoside (PGP) (5 µL of 1 mM solution for each) were spiked into each sample. Cells were lysed with FastPrep-24 homogenizer (MP Biomedical) for 30 seconds at a velocity of 4 m/s. After lysing, samples were cooled for 2 minutes on ice and then spun down for 2 minutes at 16,100 xg and 4°C. Supernatant was collected and saved after centrifugation. Cell lysis and centrifugation with the remaining sand pellet was repeated with 250 µL of MCW. After centrifugation, the supernatant was added to the previously collected supernatant for each sample. 250 µL of 80% ethanol was added to the remaining sand pellet. Samples were briefly vortexed and then incubated on a heat block at 80°C and 500 rpm for 30 minutes. After incubation, samples were spun down for 2 minutes at 16,100xg and 4°C. The ethanol supernatant was then collected and combined with the previously collected supernatant. Lysis tubes containing protein, cell debris, and the sand pellet were frozen at -70°C until protein extraction could be performed. 400 µL of water and 100 µL of chloroform were added to each of the collected supernatant tubes. Supernatant samples were then centrifuged for 5 minutes at 16,100xg and 4°C to separate phases. The upper phase (methanol, water, and polar metabolites) was collected into a new tube and dried using a step-wise pressure procedure. Dried samples were stored at -70°C until derivatization.

*Metabolite Derivatization and GC-MS Analysis*

Dried and frozen samples were allowed to acclimate to room temperature (~20 minutes). Under a fume hood, samples were dissolved in 20 µL of methoximation reagent (40 mg methoxyaminhydrochloride in 1 mL pyridine). Samples were then incubated for 90 minutes at 30°C and 900 rpm. Following incubation, 80 µL of silylation mix (1 mL N-methyl-N-trimethylsilyltrifluoroacetamid spiked with 30 µL of a mix of even-number alkanes (C10-C40)) was added to each sample and samples were incubated for 30 minutes at 37°C and 900 rpm. Following incubation, samples were centrifuged for 2 minutes at 14,000xg. Supernatant was then transferred to gas chromatography microvials and sealed with crimp caps.

Samples were analyzed in three separate runs in split-less injection mode on a Pegasus® BT GC-TOF-MS (LECO Corporation, St. Joseph, MI, USA) with a standard curve of selected metabolites for absolute quantification. Loaded volume for each sample was 1 µL. Metabolites were separated using a Rxi-5 ms column (30 m length, 0.25 mm diameter, 0.25 µm film; Restek, Centre County, PA, USA). The carrier gas as helium with a flow rate of 1 mL/min. Injection was done using a S/SSL injector with an injection temperature of 230°C. The column temperature started with 70°C for 1 minute and then followed a ramp to 340°C with at heating rate of 9°C/min. Temperature was held at 340°C for 15 minutes at the end. For the mass spectrometer, ion source and transfer line temperature was 250°C and acquisition delay was set to 300s. Masses were recorded in the range of 50-600 *m/z* with an acquisition rate of 10 spectra/sec.

Resulting peaks from selected metabolites were manually selected using LECO ChromaTOF software. Selected metabolites were absolutely quantified and normalized based on added extraction standards (see Data Analysis).

*Protein Extraction*

Sand pellets containing cellular debris and proteins were thawed on ice. Once thawed, sand pellets were washed with 500 µL of methanol. Samples were then centrifuged for 5 minutes at 16,100xg and 4°C. Methanol supernatant was then discarded and pellets were allowed to air dry under a fume hood.

To begin the extraction, 1 mL of TRIzol reagent (Thermo Fisher) was added to each pellet and mixed with the sand through gently pipetting. Samples were then incubated at room temperature for 15 minutes. Following incubation, 100 µL of chloroform was added and tubes were inverted 5 times. Samples then incubated at room temperature for three minutes. Following the incubation with chloroform, samples were centrifuged at 16,100 xg for 2 minutes at 4°C to separate phases. The lower phase, containing the proteins, was removed to a new lo-bind protein Eppendorf tube. 550 µL of water (or a 1:1 ratio with chloroform phase) was added to each sample. Tubes were inverted 5x and incubated at room temperature for 3 minutes. After incubation, tubes were centrifuged for 2 minutes at 16,100xg and 4°C. The apolar (bottom phase) was transferred to a new 2 mL low-bind protein Eppendorf tube. 1.5mL of ice cold 0.1 M $NH_4CL$ in methanol with 0.5% β-mercaptoethanol was added to each sample. Samples were then incubated on ice for 1-4 hours before being stored at -20°C overnight to facilitate protein precipitation.

*Protein Washing*

After overnight incubation at -20°C, samples were centrifuged for 15 minutes at 16,100xg and 4°C. The supernatant was discarded and 1.8 mL of ice cold methanol was added to each sample. Pellets were sonicated in an ice water bath with a Transsonic 700/H water bath (Elma) sonicator for 10-15 minutes or until pellets were completely dissolved. Samples were then centrifuged for 10 minutes at 16,100xg and 4°C. Supernatant was discarded and the washing and sonication steps were repeated with another 1.8 mL of ice cold methanol. After removal of the methanol supernatant from the second washing step, 1.8 mL of ice cold acetone was added to each sample. Pellets were again suspended in an ice water bath via sonication and then centrifuged for 15 minutes at 16,100xg and 4°C. The acetone supernatant was removed and pellets were air dried in a fume hood for 5-10 minutes (while avoiding over-drying). Once dry, proteins were stored at -70°C until further processing.

*Protein Digestion*

Dried protein pellets were resuspended in 500 µL of extraction buffer (8M urea in 50 mM HEPES, pH 7.8). Samples were then incubated for 30 minutes on a shaker at 4°C and 900 rpm. After incubation, protein concentrations were measured using the Bradford assay (Bio-Rad Cat. No. 500-0006) using a pre-made standard curve with bovine serum albumin (BSA) and an absorbance of 545 nm. The volume to get 25 µg of protein was calculated and taken for each sample and put into new low-bind protein Eppendorf tubes. Extraction buffer was added to each sample to reach a final volume of 220 µL. 5.64 µL of 200 mM dithiothreitol (DTT) was added to each sample and samples were then incubated for 45 minutes at 37°C and 700 rpm. Next, 2.26 of freshly prepared 1 M of iodoacetamide (IAA) was added and samples were incubated for 60 minutes at 30°C and 700 rpm. After the IAA incubation, a final amount of 6.27 µL of 200 mM DTT was added and samples were again incubated in the dark at room temperature for 15 minutes. Next 5 µL of 0.1 µg/µL of mass spec grade rLysC (Promega) was added to each sample and proteins were digested for 3 hours at 37°C and 700 rpm. Following the rLysC digestion, 660 µL of trypsin buffer (2 mM $CaCl_2$, 5 mM DTT, 50 mM $NH_4HCO_3$, 10% acetonitrile (ACN)) was added to each sample. After the addition of trypsin buffer, 1 µL of trypsin beads from a Poroszyme Immobilized Trypsin Cartridge (Applied Biosystems) were added to each sample. Samples were incubated with trypsin beads for 16 hours at 30°C on a rotator. After digestion, samples were stored at 4°C (briefly, not all can be desalted at once) until they could be desalted. Following digestion, peptides were desalted using OMIX C18 pipette tips (Agilent Technologies). For this, samples were centrifuged briefly to pellet the trypsin beads before desalting. Tips were activated by washing with 100 µL of methanol and then washed twice with 100 µL of 0.1 % formic acid (FA). Peptide solution was then acidified with the addition of FA to a final concentration of 3 %. The solution containing peptides was pipetted through the C18 tips and then washed twice with 100 µL of 0.1 % FA. Peptides were then eluted and saved from the C18 tips by being washed twice with 100 µL of methanol. After desalting, peptides were dried in a ScanSpeed 40 speed vacuum with a ScanVac vacuum control and stored at -20 °C.

*LC-MS/MS*

To prepare the peptides for mass spectrometry analysis, peptides were resuspended in 250 µL of 2 % acetonitrile (ACN) and 0.1 % formic acid (FA). 5 µL were then injected into an EASY-Spray C18 column (2µm, 100Å 75 µm x 50 µm, Thermo). Peptides were eluted from the column for 150 min using a 90 min linear gradient starting from 96 % solvent A (0.1 % FA) and 4 % solvent B (80 % ACN, 0.1 % FA) to 35 % of solvent B with a flow rate of 0.3 µL/min. The linear gradient was followed by an increase of solvent B to 90 % over 1 min and held at 90 % for 8 min. Solvent B was then adjusted from 90 % to 4 % over 1 min and held at 4 % for the remaining 50 min. Ion source was an EASY-Spray source with a spray voltage of 1.9 kV. Mass spectrometry measurements were taken using an LC-QExactive-Plus (Thermo) with the following settings: 0 to 150 min; MS1: positive polarity, full scan range 380-1800 *m/z*, resolution 70,000, collision-induced dissociation (CID) fragmentation for the 20 most intense ions; MS2: loop count 20, resolution 17,500, scan range 200-2000 *m/z.*

#### **Data Analysis**

*DNA Production, Protein Production, and Carbon Consumption*

Values for DNA (µg), protein (µg), and consumed carbon (mmol) were normalized to the total amount of ammonia/ammonium consumed in each culture (mmol).  Normalized values were subjected to a box plot analysis as described below for protein and metabolite values. For carbon consumption some samples were removed:A1-A3 did not have GC measurements available; E2, G1, and G2 as carbon consumption values were either negative (E2) or too low to be considered biologically accurate (G1 and G), likely due to sampling errors when collecting the gas phase.

*Metabolites*

Extracted metabolites were randomly distributed between three batches to be run on the GC-MS. Quality control samples containing known concentrations of metabolites were run for each batch individually. Detected trimethylsilyl (TMS) groups were combined for each metabolite.  The internal standard PGP was used to normalize sample as it showed less variation than the PE internal standard. Based on an initial analysis of metabolite areas, some contamination could be seen from blank samples that contained only purified water. Therefore, PGP normalized areas from blank samples were subtracted from samples within their perspective runs. PGP normalized and blank corrected sample values were then used to determine absolute quantification of metabolites.

Standard curves were manually created for each metabolite in each batch. In cases where concentrations on the standard curve became non-linear, only quality curve points that captured the spread of areas from samples were used (i.e. if only low concentrations were detected in samples, high values of standards were disregarded). Linear regression was applied to selected quality control points and slope and intercept values were used to obtain pmol values for metabolites.

Absolute amounts of metabolites for each culture were converted into carbon-moles (C-Moles) and used in conjunction with the amount of carbon consumed for each culture to normalize values to show how much consumed carbon was allocated for each metabolite (i.e. C-mol metabolite per C-mol carbon consumed). Carbon normalized values were subjected to a boxplot analysis for each metabolite in each condition. Based on this analysis outliers were removed. Normalized values with outliers removed were then hierarchically clustered using the heatmap.2[8] function in R version 4.3.1[9].

*Proteins*

Protein identification from raw mass spectrometry data was analyzed using MaxQuant version 1.6.6.0[10]. Max missed cleavages was set to 2 with variable modifications of "Oxidation (M)" and "Acetyl (Protein N-term)", fixed modifications set to "Carbamidomethyl (C)", an FDR of 0.01, and the digestive enzyme as Trypsin/P and LysC. Minimum number of unique and razor peptides was set to 2 and protein values were LFQ normalized. Reference proteome for *N. viennensis* was downloaded from Uniprot in spring of 2019.

LFQ normalized data was analyzed using a boxplot analysis for each protein in each condition. Identified outliers were removed and averages were calculated for each protein in each group. A PCA plot of the cleaned up data was made using the fviz_pca_ind function of the factoextra package[11] in R. Cleaned up and averaged data were hierarchically clustered using the heatmap.2 [8] function in R and split into 7 clusters. For mean comparisons, individual proteins were tested for normality (Shapiro test, shapiro_test() function of rstatix package[12]) and homogeneity of variance (Levene test, leveneTest() function of car package[13]). If all conditions passed these tests, one-way ANOVA (aov() function, stats package[9]) was used to determine if protein averages varied among conditions. If all conditions did not pass these tests, a Kruskal-Wallis test (Kruskal.test() function, stats package[9]) was used to determine if protein averages varied among conditions. For proteins of interest, Tukey tests (TukeyHSD() function, stats package[9]) or Dunn's test (dunnTest() function, FSA package[14]) was used to determine which conditions were different for proteins with ANOVA and Kruskal-Wallis analysis respectively. In each case, *P* values were adjusted using the Benjamini-Hochberg method (p.adjust() function, method="BH", stats package[9]).

Clusters of interest for the most limited condition (E; Figure 3, Cluster V ) and non-limited conditions (ACDG; Figure 4, Cluster VII) were identified by choosing the clusters with the highest average values for condition E and ACD respectively.

Proteins were evaluated based on functional categories according to their archaeal clusters of orthologous groups (arCOG) categories as defined in Reyes et al. (2020)[15]. For arCOG enrichment analysis, the phyper() function of the stats package[9] was used for defined clusters or groups of proteins and compared against all detected proteins in the dataset with the parameter lower.tail=FALSE. *P* values for each group of proteins were adjusted using the Benjamini-Hochberg method using the p.adjust() function of the stats R package[9].

*Statistical Data Analysis*

Clusters of high relative abundance for metabolites in condition E (Figure 5, Cluster Met-IV) were correlated with all proteins across all conditions to identify trends in metabolism. The R function cor.test was used with method set to "pearson" and use set to "complete.obs". *P* values were adjusted according to the Benjamini-Hochberg method.

A correlation analysis was performed with all proteins for all conditions against MsrA to identify other proteins with similar patterns. The R function cor.test was used with method set to "pearson" and use set to "complete.obs". *P* values were adjusted according to the Benjamini-Hochberg method.

Correlation analyses for specific metabolites (glucose, trehalose, maltose, and melibiose) were performed across all proteins and conditions. The R function cor.test was used with method set to "pearson" and use set to "complete.obs". *P* values were adjusted according to the Benjamini-Hochberg method.

Other R packages used in analysis included ggplot2[16], dplyr[17], reshape2[18], tidyverse[19], ggVennDiagram[20], RColorBrewer[21], expss[22], magrittr[23], plot.matrix[24], and viridis[25].

*Partial Least Squres-Discriminant Analysis (PLS-DA)*

A PLS-DA analysis was performed with all proteins to identify specific proteins of interest under carbon limitation. Outlier protein values were replaced with averages of the respective protein for the respective condition. Protein data was then scaled using the pareto method (done for each protein across all conditions). The mixOmics R package[26] was used with the plsda command (X as the scaled proteins, Y a vector representing each condition, scale=FALSE) to create a PLS-DA plot. The function vip() was used to identify VIP (variable importance in projection) scores for proteins pertaining to component 1 which separated samples by carbon limitation. The protein with the highest VIP score in component 1 was AOA060HNZ6 (NVIE_010650), a hypothetical protein.

*Identification of NVIE_010650*

A blast search of NVIE_010650 revealed no hits with putative functions. A structural search using the AlphaFold generated structure was performed using Foldseek[27]. The hits with putative function were summarized (Supp. Data). The structures of the retrieved sequences together with that of NVIE_10650 were aligned using PROMALS3D[28]. The alignment was then filtered with TrimalAL[29] and trees were constructed with IQ-TREE v2.3.6[30] and visualized in iTOL v.6[31].

### Supplementary Figures

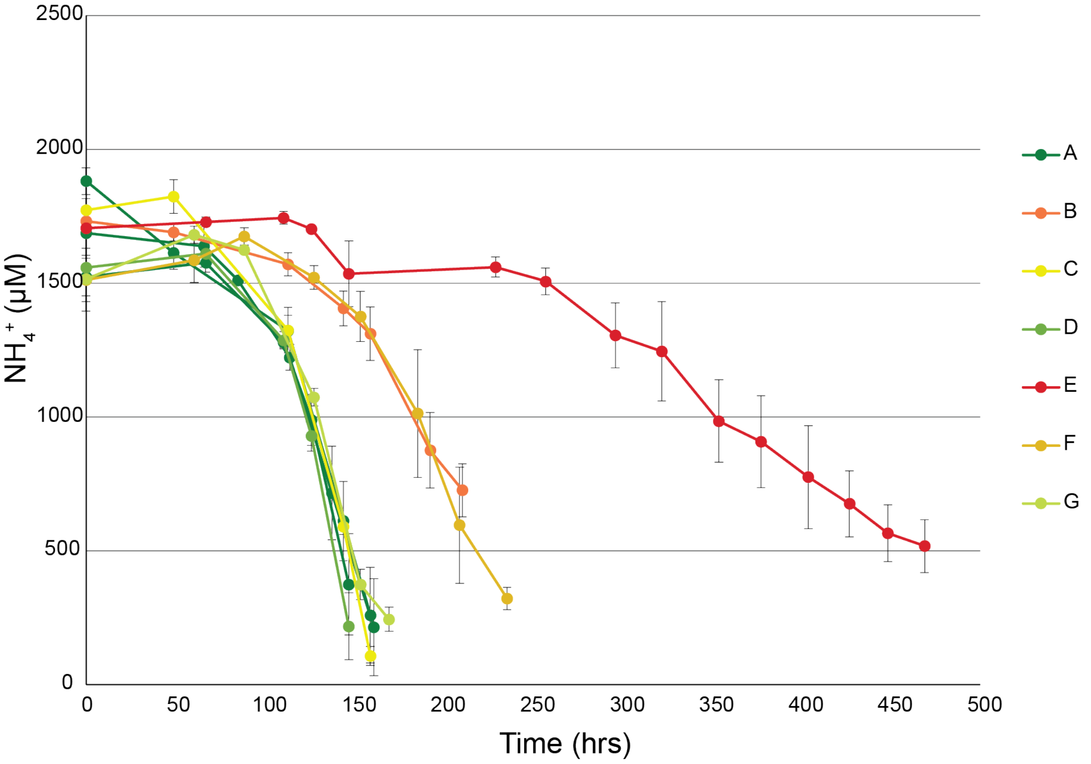


**Figure EV1: Consumption of ammonium during growth.** Error bars represent standard deviations of biological replicates.

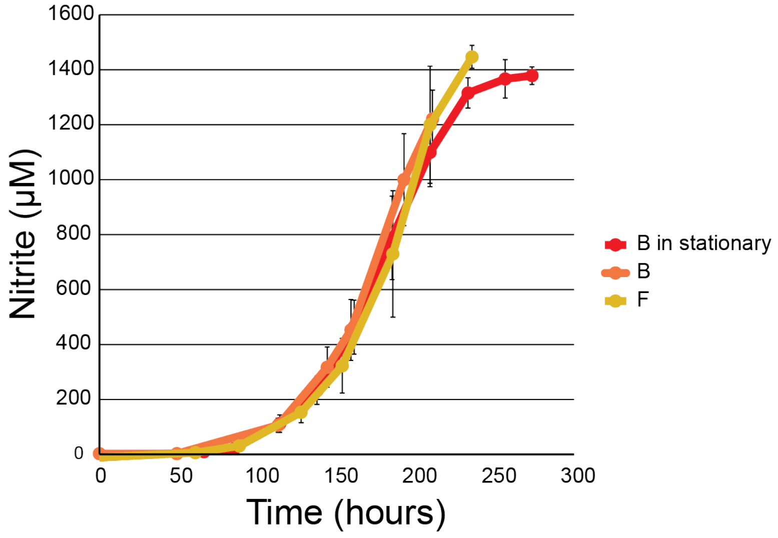


**Figure EV2: Growth curves of conditions B and F.** Growth curves showing condition F (0.1 mM carbon with pyruvate) and condition B (0.1 mM carbon with catalase). "B in stationary" represents condition B when allowed to grow above ~1200-1300 mM nitrite. At this point, the cultures begin to enter stationary phase. Condition F is able to oxidize more ammonia without entering stationary phase.

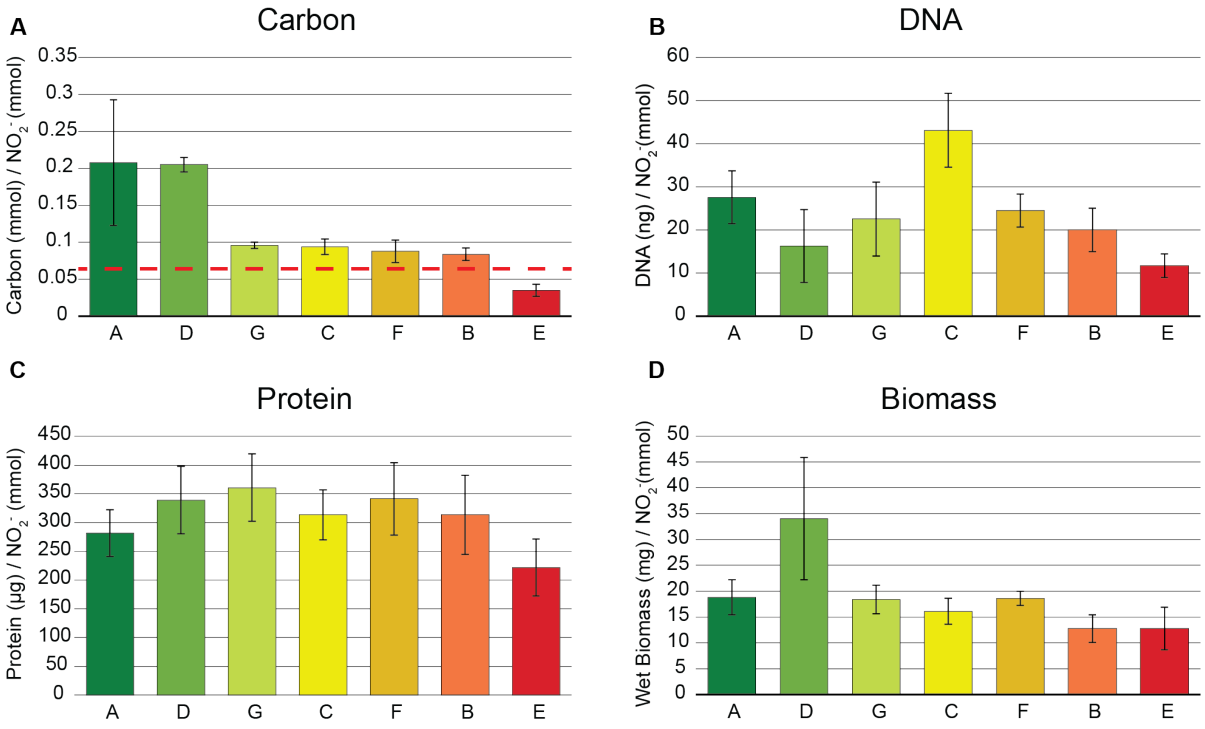


**Figure EV3: Carbon and biomass data of *N. viennensis*. A.)** Average inorganic carbon consumption normalized to nitrite production. The red dotted line shows the theoretical amount of carbon consumption for 1 mmol of ammonia consumed based on Equation 18. **B.)** Average DNA content normalized to nitrite produced. **C.)** Average protein content normalized to nitrite production. **D.)** Average wet biomass produced normalized to nitrite production. Error bars represent standard deviations of cultures.

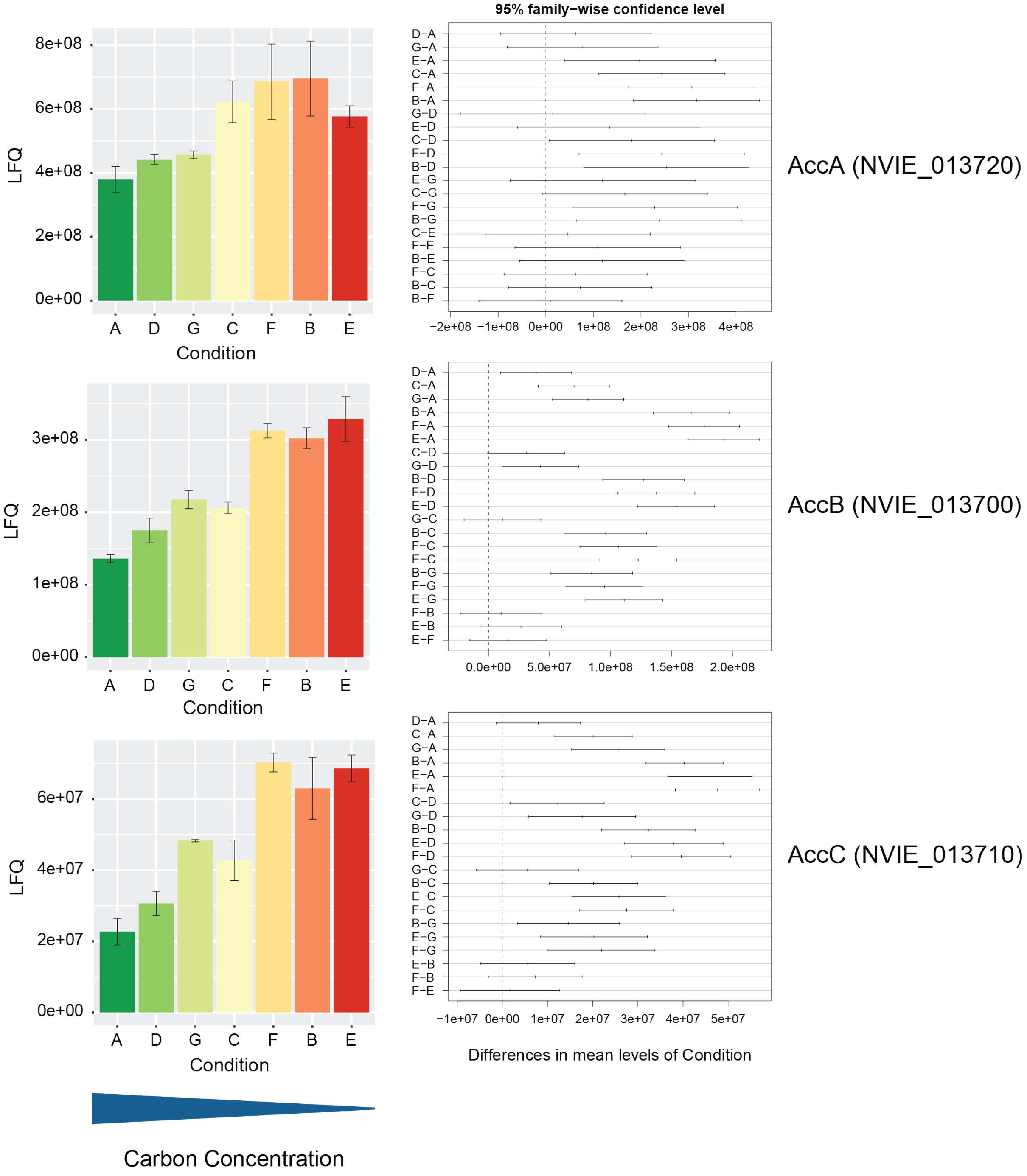


**Figure EV4: Relative intensities and 95% confidence intervals of subunits of acetyl-CoA carboxylase across conditions.** Average intensities are shown as normalized label free quantification (LFQ) values. Error bars represent standard deviations. Family-wise confidence intervals represent differences between compared conditions. Confidence intervals that do not cross 0 represent statistically significant differences between the two tested groups.

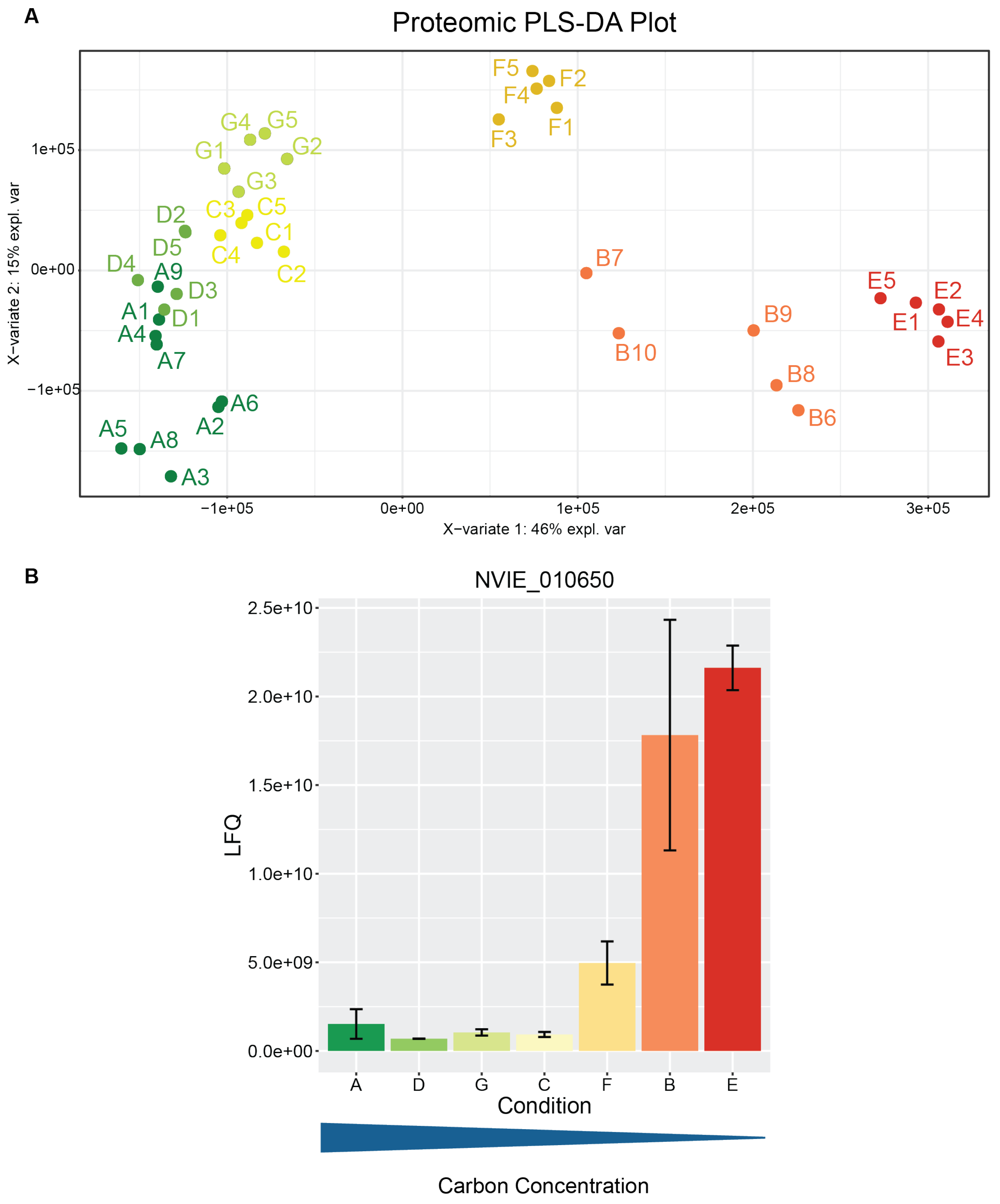


**Figure EV5: PLS-DA analysis. A.)** Plot of a supervised partial least squares discriminant analysis. **B.)** Average intensities of the protein with the strongest VIP value corresponding with X-variate 1 of the PLS-DA analysis. Average intensities are shown as normalized label free quantification (LFQ) values. Error bars represent standard deviations.

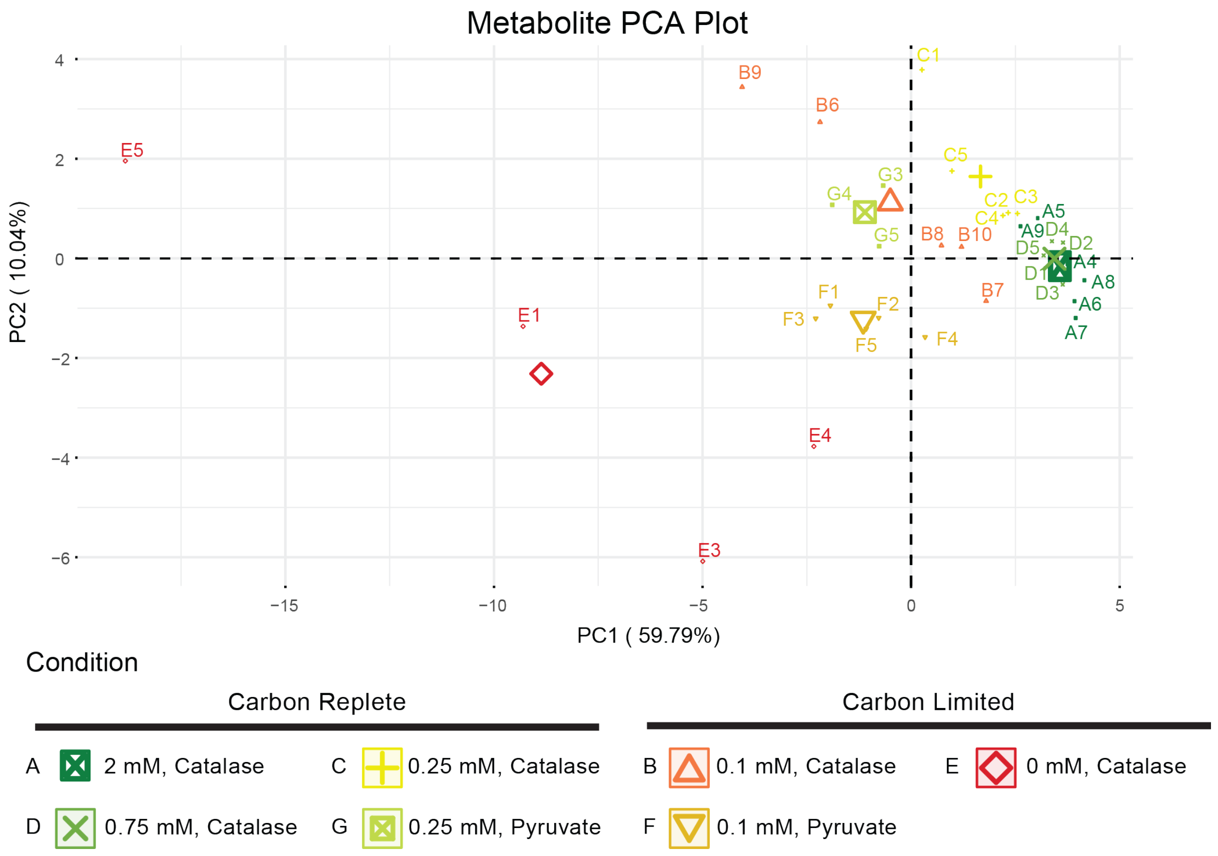


**Figure EV6: Principal component analysis of *N. viennensis* metabolomes.** Bold and enlarged points represent centers of points within respective conditions.

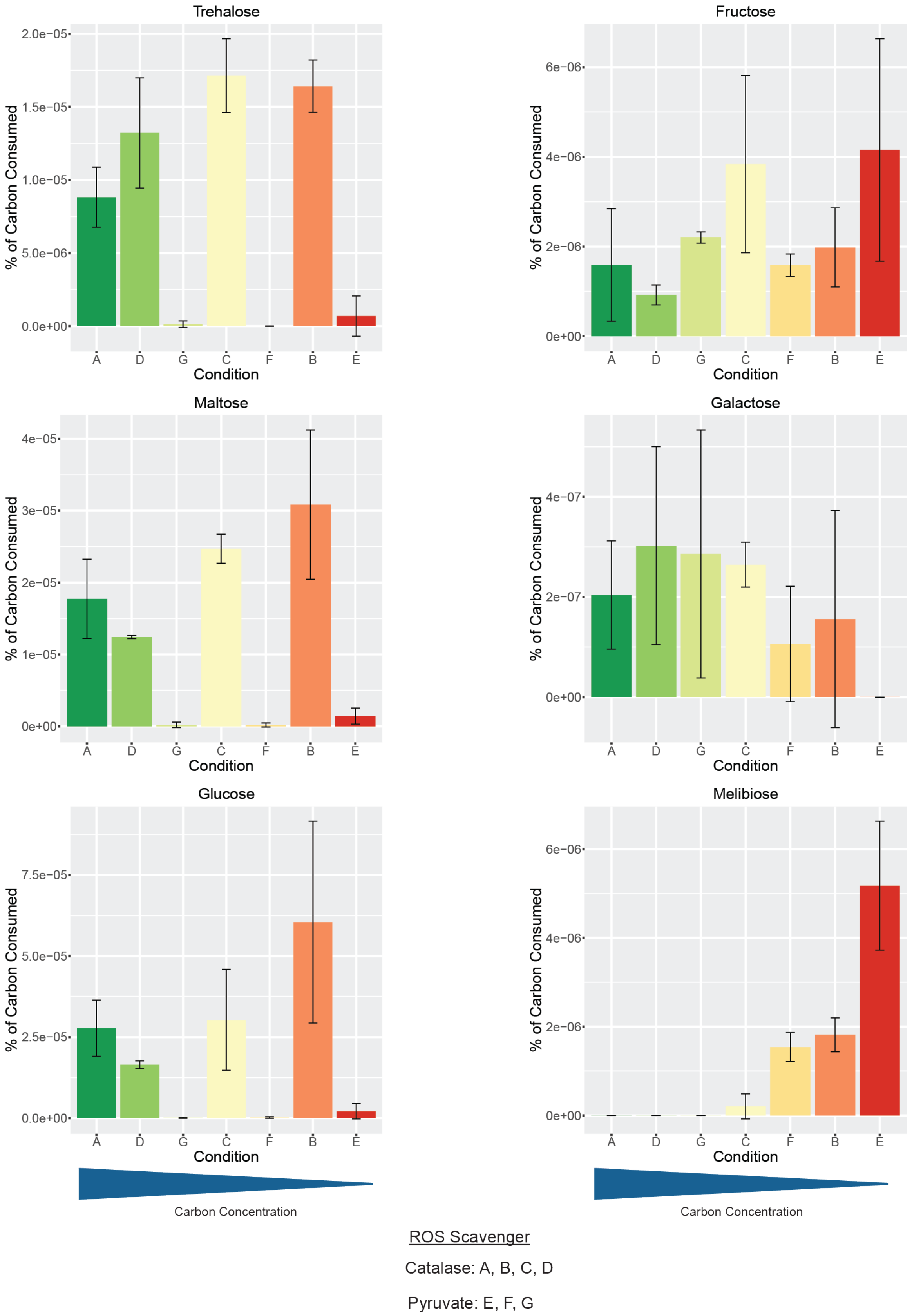


**Figure EV7: Sugar abundance across carbon conditions.** Average abundance values of quantified sugars normalized to amount of carbon consumed in each culture. Error bars represent standard deviations.

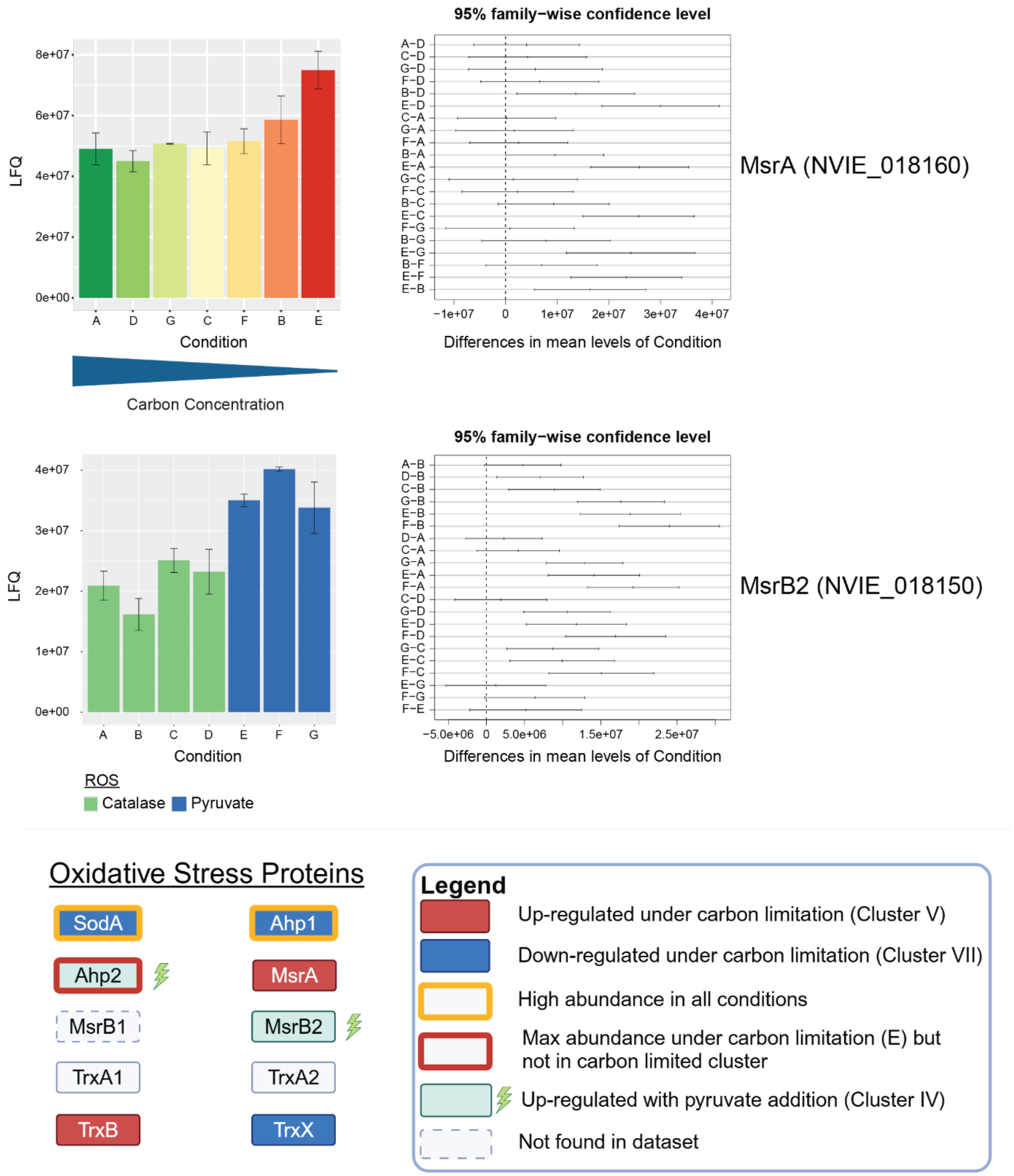


**Figure EV8: Oxygen detoxification proteins in *N. viennensis*.** Average intensities are shown as normalized label free quantification (LFQ) values. Error bars represent standard deviations. Family-wise confidence intervals represent differences between compared conditions. Confidence intervals that do not cross 0 represent statistically significant differences between the two tested groups. Locus tags and accession numbers can be found in Dataset EV1. Some pieces created in BioRender. Hodgskiss, L. (2025) https://BioRender.com/u08b790.

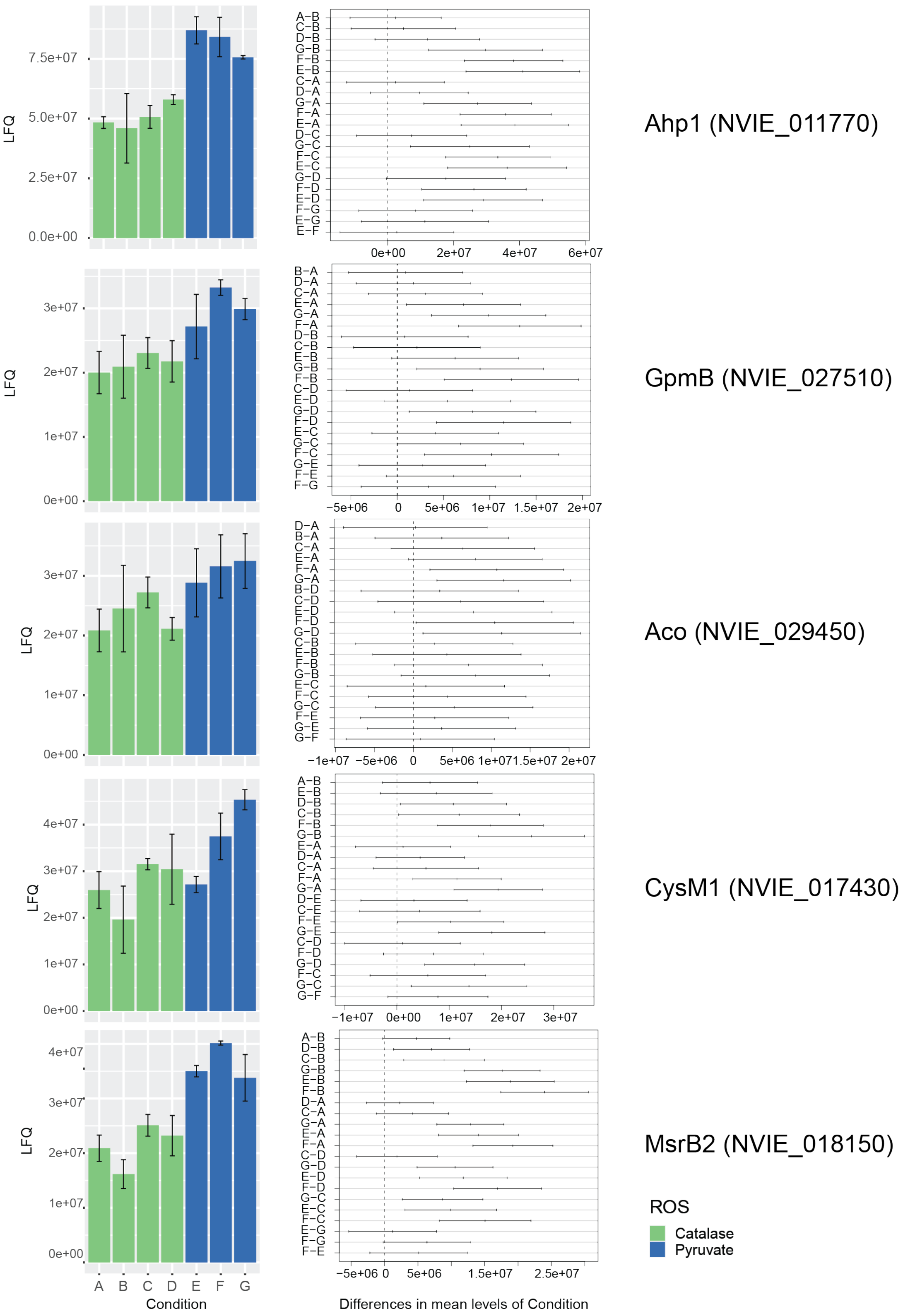


**Figure EV9: Central carbon and oxygen detoxification proteins that react to different ROS scavengers.** Average intensities are shown as normalized label free quantification (LFQ) values. Error bars represent standard deviations. Family-wise confidence intervals represent differences between compared conditions. Confidence intervals that do not cross 0 represent statistically significant differences between the two tested groups.

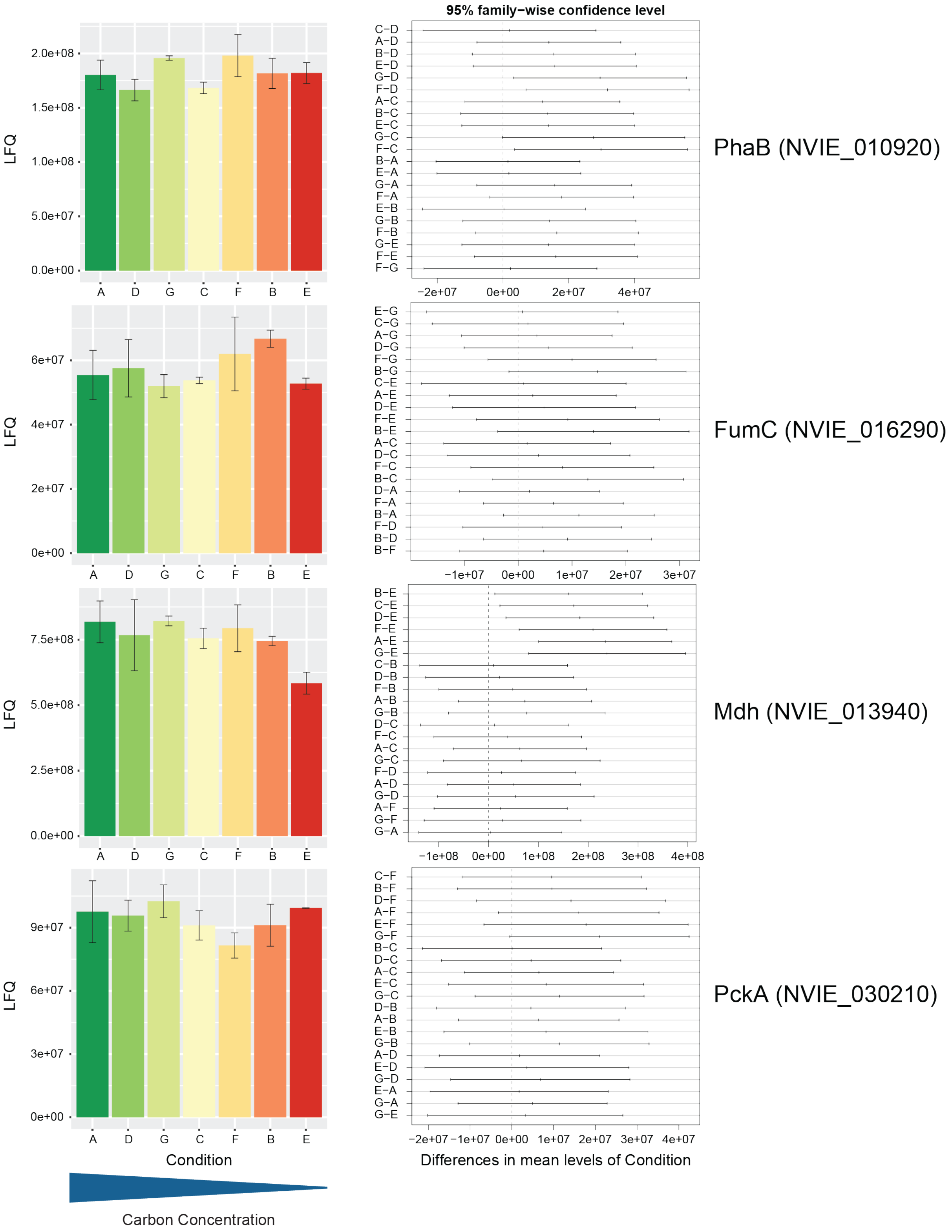


**Figure EV10: Proteins of the central carbon metabolism that do not increase under carbon limitation.** Average intensities are shown as normalized label free quantification (LFQ) values. Error bars represent standard deviations. Family-wise confidence intervals represent differences between compared conditions. Confidence intervals that do not cross 0 represent statistically significant differences between the two tested groups.

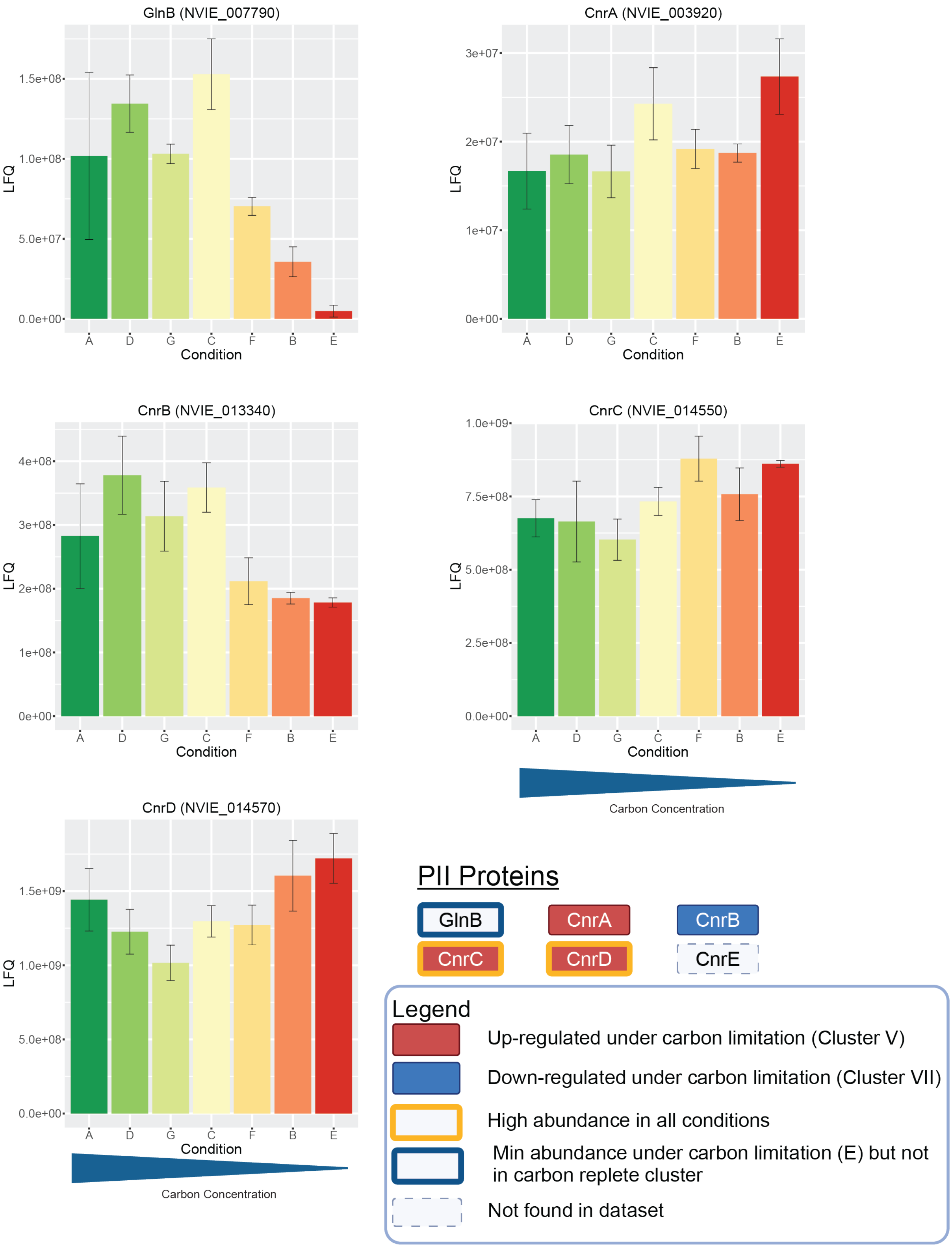


**Figure EV11: PII protein response to carbon limitation.** Average intensities are shown as normalized label free quantification (LFQ) values. Error bars represent standard deviations. Family-wise confidence intervals represent differences between compared conditions. Confidence intervals that do not cross 0 represent statistically significant differences between the two tested groups. Some pieces created in BioRender. Hodgskiss, L. (2025) https://BioRender.com/j28f691.

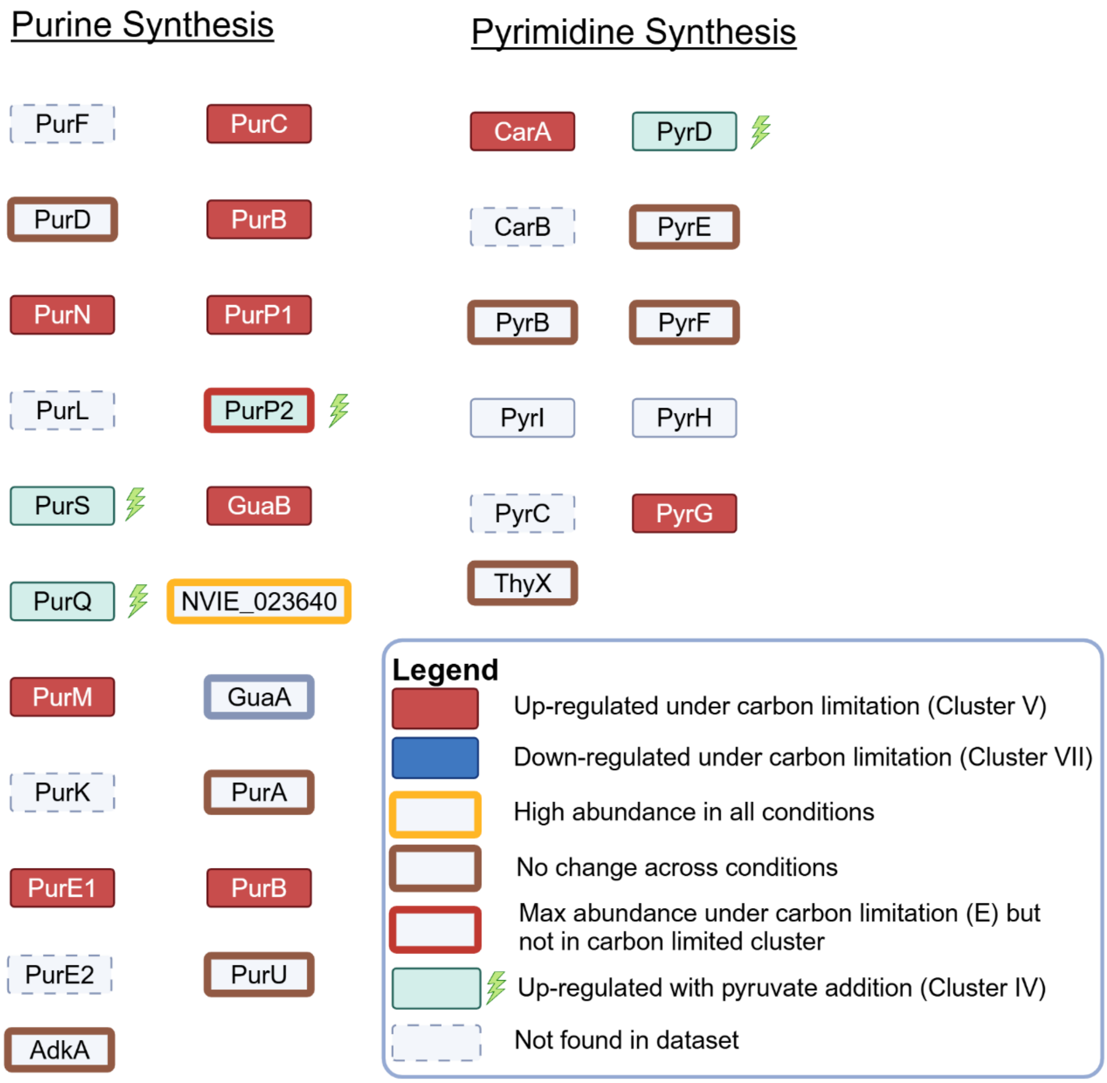


**Figure EV12: Purine and pyrimidine synthesis proteins in *N. viennensis*.** Locus tags and accession numbers can be found in Dataset EV1. Created in BioRender. Hodgskiss, L. (2025) https://BioRender.com/c49z928.

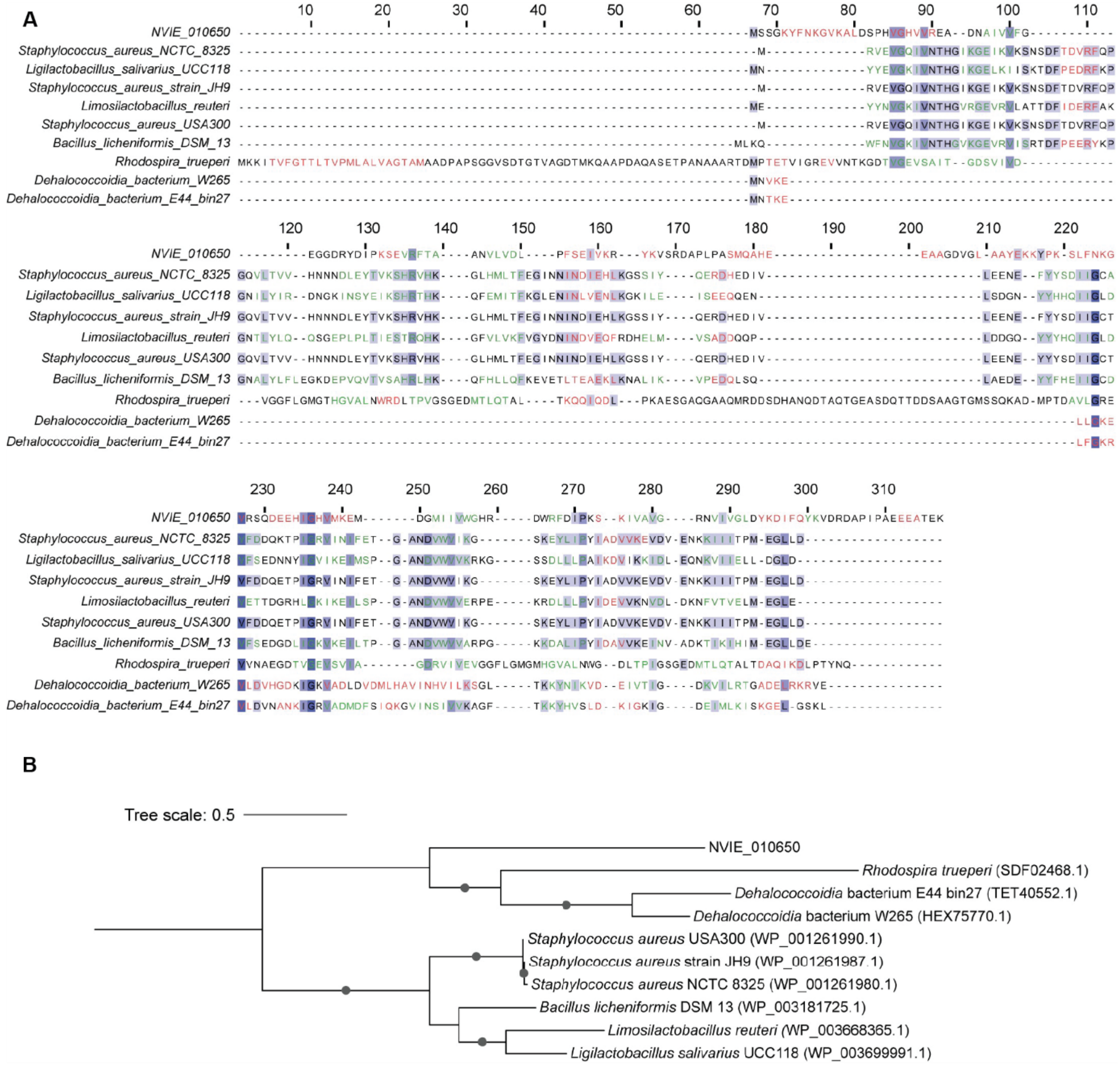


**Figure EV13: Analysis of structural search matches for NVIE_010650. A.)** Amino acid alignments of NVIE_010650 with closest structural search matches from FoldSeek. **B.)** Amino acid tree showing the relatedness of NVIE_010650 with closest structural matches from FoldSeek.

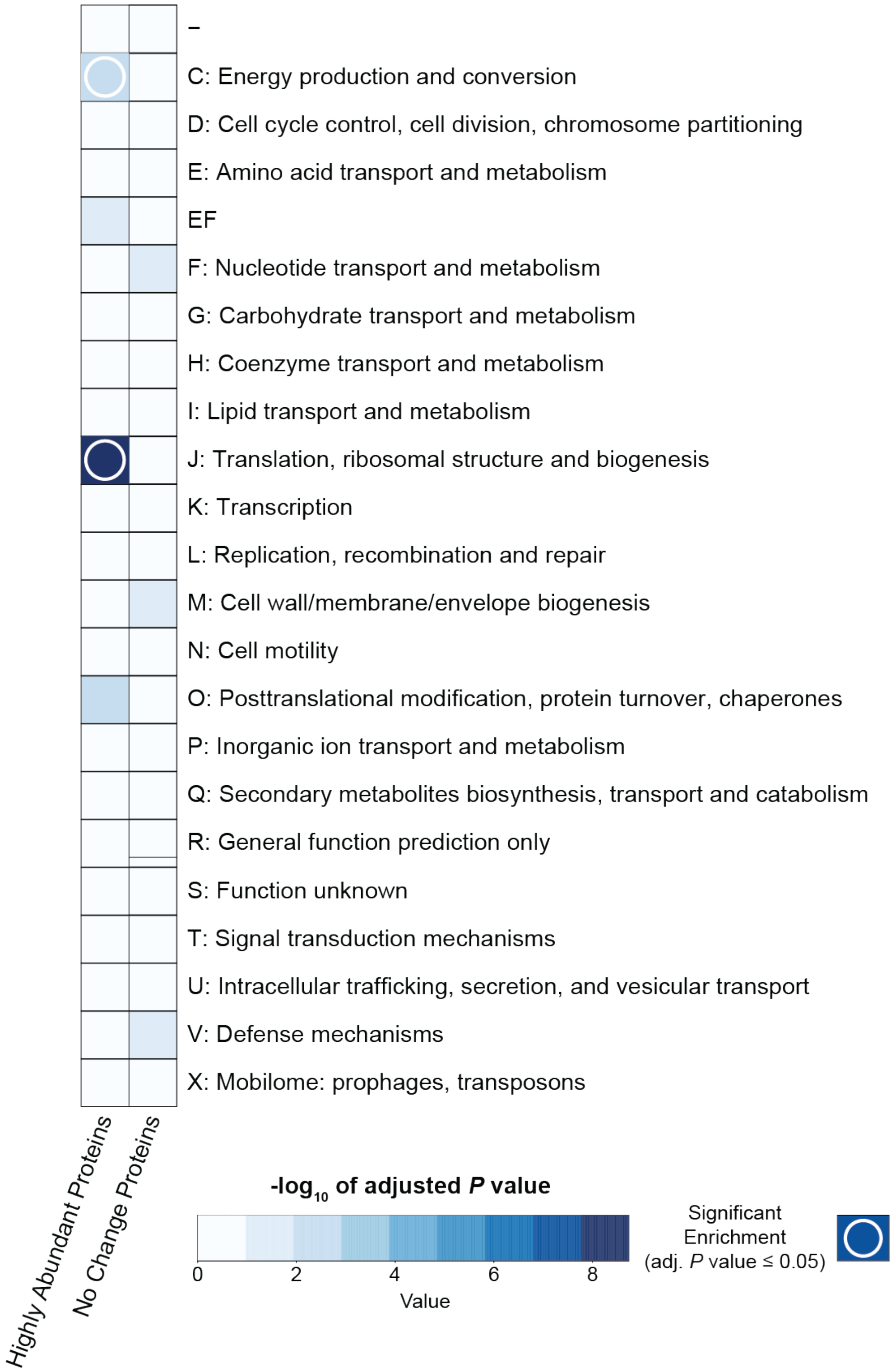


**Figure EV14: Enrichment analysis of arCOG categories in highly abundant proteins and proteins that do not change between different conditions.** Boxes with a white circle indicate arCOG categories that are enriched in their respective cluster based off of a hypergeometric test.
